# SCDT: Detecting somatic CNVs of low chimeric ratio in cf-DNA

**DOI:** 10.1101/411256

**Authors:** Zhaoyang Qian, Xiaofeng Wang, Chang Shi, Rui Han, Yaoshen Wang, Hongmei Zhu

## Abstract

**Motivation:** Sequencing of cell-free DNA (cf-DNA) has enabled Noninvasive Prenatal Testing (NIPT) and”liquid biopsy” of cancers. However, while the aneuploidy and point mutations were focused on by most of NITP and liquid biopsy studies, detecting sub-chromosome CNVs that affect a few to dozens of megabases was rarely reported, likely attributable to the difficulty in accurately identifying them, especially for those present in a small fraction of cf-DNA.

**Results:** We developed a somatic CNV detection tool (SCDT), for detecting sub-chromosome CNVs in cf-DNA using whole genome sequencing (WGS) data or off-target reads in target sequencing data. Additional to using control samples for correcting genome position specific bias, two GC correction steps were performed, which regressed GC content of DNA fragments and that of genome bins, respectively. After GC correction, the coefficients of variation of copy ratios approximated the lower boundary of theoretical values, suggesting removing of almost all systematic errors. Finally, CNVs were detected by a piecewise least squares fitting based segmentation algorithm, which outperformed other segmentation methods. We applied SCDT on simulated and real maternal plasma samples, and target cf-DNA sequencing of 118 normal individuals and 240 cancer patients, and demonstrated high sensitivity and specificity.

**Availability:** SCDT is available at https://github.com/Martiantian/Somatic_cnv_detect_tool.

**Supplementary Information:** Supplementary data are available at Bioinformatics online

## 1 Introduction

Copy number variation (CNV) is one type of structural variation with duplication or deletion event that affects a considerable number of base pairs (Sharp, *et al*., 2005), altering gene dosage and subsequently affecting functional and biological behavior of cells. CNVs have been known to greatly contribute to a wide repertoire of human diseases, including genetic disease, developmental and neuropsychiatric disorders (Kirov, *et al*., 2009; Sebat, *et al*., 2007; Walsh, *et al*., 2008) and almost all types of cancers (Pollack, *et al*., 2002; Shlien and Malkin, 2009; Taylor, *et al*., 2008). Detecting CNVs in human genomes has been a routine clinical test for disease screening, diagnosis and therapy guiding.

In the recent years, plasma cell free DNA (cf-DNA) sequencing has been broadly applied to non-invasive genetic diagnostics. One of the most important applications of cf-DNA sequencing is non-invasive prenatal testing (NIPT), which directly sequences cf-DNA extracted from maternal blood to identify likely aneuploidy of the fetus. However, NIPT generally focused on whole-chromosome aneuploidies (triploid 13/18/21, X, XXY and XYY) which account for only 30% of all live births with a chromosome abnormality. Recent progression has been made in genome-wide screening of sub-chromosomal CNVs with significantly smaller sizes, which have a considerably higher incidence (0.5-1.7%) than whole-chromosome aneuploidy in human pregnancy (Brady, *et al*., 2016), and could be associated with genetic disease including DiGeorge syndrome (22q11.2 deletion), Cri-du-chat syndrome (5p deletion), Angelman syndrome (15q11–q13 deletion) and 1p36 deletion syndrome. However, non-invasively detecting CNVs with small chimeric fraction without previously known positions is much more challenging than aneuploidy testing. After two proof-of-concept studies (Jensen, *et al*., 2012; Peters, *et al*., 2011) on a few cases, several subsequent studies focused on achieving statistical significance using relatively high depth whole genome sequencing (more than 100 million reads). These methods were based on statistical test on individual genomic bins, or required several consecutive bins to be significant (Srinivasan, *et al*., 2013; Yu, *et al*., 2013) However, in addition to the cost of relatively deep whole genome sequencing, high rates of false positives (FPs) and false negatives (FNs) in these individual-bin test methods would restrict their application in real clinical use. Some other methods reduced the requirement on sequencing depth by employing sliding window strategy (Straver, *et al*., 2014) or using binary segmentation with dynamic threshold (Chen, *et al*., 2013), and aimed at detecting large fragment aberrations (>10M) using low-coverage sequencing data. Lo et al, reported 60.7% (17/28) of accuracy for analyzing 3Mb to 42Mb de novo CNVs using 4-10 M reads, while the sensitivity increased to 92.9% using relative higher sequencing depth up to 120M reads (Lo, *et al*., 2016). Yin et al developed a method to identify 69 of 73 (94.5%) CNVs identified by array CGH using 10 million reads, with a specificity of 98.1% (Yin, *et al*., 2015). A method based on unified Hidden Markov model was developed for detecting fetal CNVs and achieved great resolution (400 kb) with fetal fraction of 13%, however, is only feasible using deep sequencing data and information of parental SNP genotypes, which is unavailable in routine NIPT (Rampasek, *et al*., 2014).

Another inspiring application of cf-DNA sequencing is to be used as a surrogate for tissue biopsy, named”liquid biopsy”, for screening and monitoring tumor-derived genomic aberrations. Circulating tumor DNA (ct-DNA) can be detected in the plasma of cancer patients, and has great potential in clinical management of cancers. A plenty of targeted therapies have been developed to target copy number change of some cancer driver genes, e.g. high level amplifications of ERBB2, MET, CCND1 and FGFR1 (Baselga and Swain, 2009; Christensen, *et al*., 2005; Musgrove, *et al*., 2011; Turner, *et al*., 2010), etc. However, current reported noninvasive assays seldom include detection of actionable CNVs, which may be due to the serious difficulty on accurately identifying CNVs in ct-DNA, as ct-DNA typically accounts for a little proportion (<10%) of cf-DNA, even in many advanced stage cancer patients(Adalsteinsson, *et al*., 2017). Most published studies employed reads counting strategy for genome bins and simple statistics such as Z-test to identify CNVs. Chan et al used individual bin based Z-test on 4 high depth whole genome sequencing (WGS) (17X) of HCC cases (Chan, *et al*., 2013). Heitzer et al calculated segment z-score after CNV segmentation using circular binary segmentation (CBS) algorithm in 13 plasma samples of 9 metastatic prostate patients, which usually have high tumor DNA concentrations (Heitzer, *et al*., 2013). Xu et al performed individual bin z-score based CNV analysis on 31 patients, and showed that recognizable CNVs were only detectable in most samples with large tumor size (tumor dimension > 50 mm) (Xu, *et al*., 2015). However, method accurately determining the CNV fragment using shallow depth WGS data with low FPs is scarce and tools for identifying tumor-derived CNV in cf-DNA of a wide range of patients would have significant clinical values.

In this study, we present a novel approach named Somatic CNV Detecting Tool (SCDT), which has ability to use shallow WGS data and target sequencing data to detect genome-wide microdeletions or microduplications (MDs) without a priori knowledge of an event’s location. To maximize the ability to remove “noise” introduced by library construction, PCR process and sequencing, and intrinsic difference between genome regions, we used control samples to correct genome position specific bias, and two GC correction steps to regress GC content of DNA fragments and that of genome bins, respectively. Following that we nearly achieve the theoretical minimum of random fluctuation in copy ratios. A segmentation algorithm based on piecewise least squares fitting and a rigorous statistical method is applied to finally determine the CNVs. We show that SCDT recovered all “spiking in” MDs with ≥3Mbp length and ≥5% chimeric fraction (≥10% mix ratio) using about 25 million sequencing reads (0.3× genome coverage). Finally, we applied our algorithm in three cf-DNA datasets of abnormal maternal plasma samples, normal samples and a large cohort of patients with various types of cancer, respectively, and demonstrated the feasibility of SCDT for precisely detecting clinically relevant CNVs in cf-DNA.

## 2 Methods

### 2.1 Data preparation and Overview of methods

We use bam format files as input of SCDT, including at least one sample as control. The control samples should be prepared using the same protocol with the test samples, including methods for library constructing and sequencing. Duplicate reads should be removed in the bam files using software such as Picard-tools and Samtools. The aligned genome positions of reads were extracted from the bam files, with a filtering step to discard reads with low mapping quality or high number of mismatches. Additionally, for using target sequencing data as input, reads located adjacent to (<500 bp) the target region were discarded from the bam files.

Firstly, the whole genome should be divided into non-overlapping bins (defined as level-1 bins) with fixed length assigned by users. Read depth count (RDC) of each level-1 bin is obtained by counting reads with start positions in it. However, each DNA fragment was not counted by 1, but instead by 1 divided by a correction factor corresponding to its CG content (section 2.2). Secondly, the RDCs of each test and control sample were centralized to 1 by dividing their medians, and then we used the mean of centralized RDCs at each bin across the control samples to generate a reference data, which is used to normalize the centralized RDCs of test samples to obtain the copy ratios (section 2.3). Thirdly, we merged a fixed number (defined by users) of level-1bins into the level-2 bin. The copy ratio of each level-2 bin was calculated as the mean of copy ratios of level-1 bins inside it. Subsequently, we performed a second step of GC correction based on regression for the copy ratios and GC content of level-2 bins using general liner model (GLM) (section 2.4). Finally, we performed CNV segmentation by a piecewise least squares fitting on the copy ratios of level-2 bins (section 2.5), and tested the significance of each CNV segment under the assumption of independent and identical distribution of copy ratios (section 2.6) (Supplementary Fig S1).

### 2.2 First step of GC correction based on single DNA fragment

Firstly, we divided the GC-content range (0-1) into 1000 intervals (0.001 per interval), followed by calculating GC-content distribution on the 1000 GC intervals for 170 base-pair (bp) sliding windows (sliding with 1bp each time) in the reference genome (except the sex chromosomes). Secondly, for single-end sequencing data, we extended the sequencing reads to 170bp based on its alignment position in the reference genome. Then we calculated GC-content distribution of extended reads on the 1000 GC intervals for each sample. GC-content intervals higher than 70% and lower than 20% were discarded for their intense fluctuation of read counts. For each DNA fragment, a correction factor *cf* was assigned by:

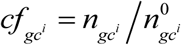

Where *gc*^*i*^ is GC interval of the DNA fragment i, 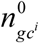 is distribution density of *gc*^*i*^ in the reference genome, 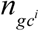 is distribution density of *gc*^*i*^ in sequenced DNA fragments of this sample (Fig 1A). So we can get the normalized read depth count (NRDC) of each level-1 bin by

**Figure 1.**
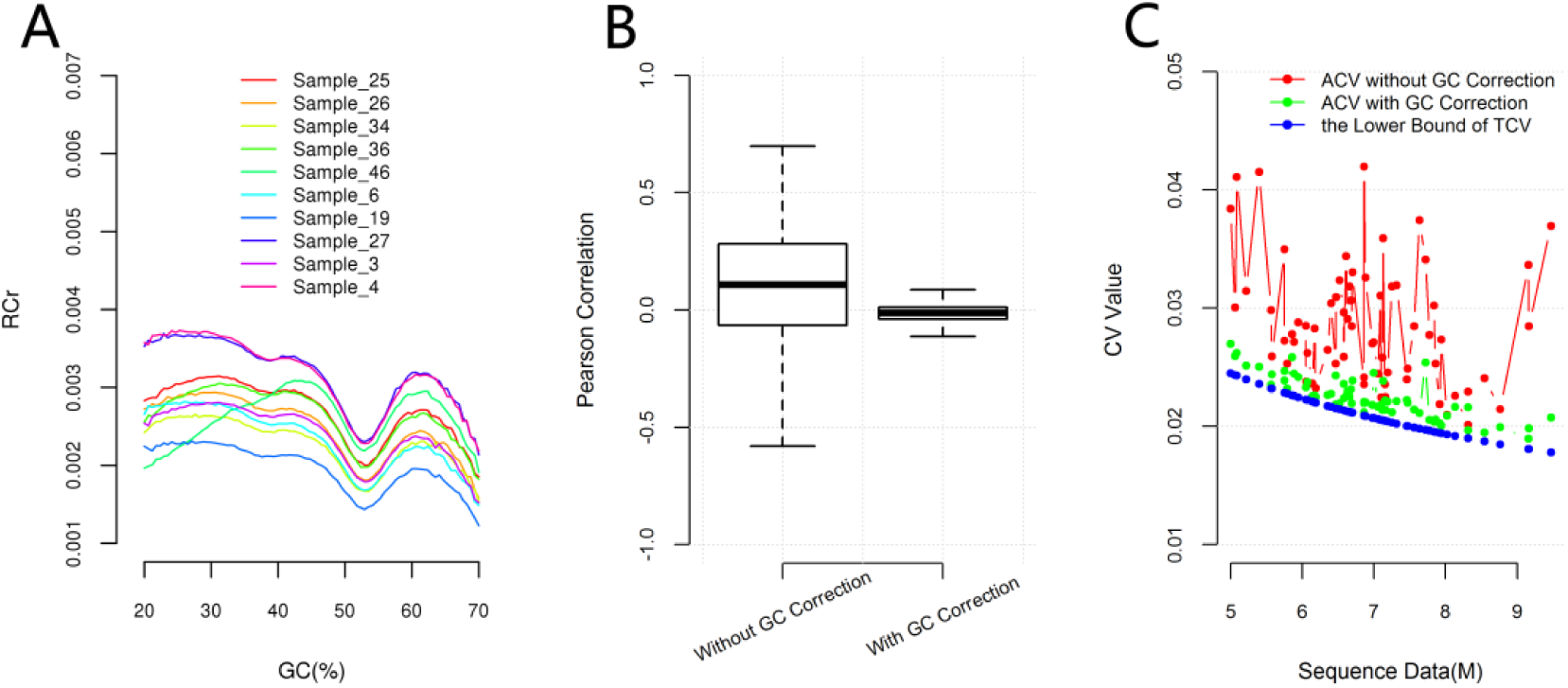
GC correction in 67 normal maternal plasma samples. (**A**) GC correction factors for DNA fragments based on the GC content by equation (2). This figure only shows ten samples randomly selected from the 67 normal maternal plasma samples. (**B**) Comparison of Pearson’s correlation between copy ratios of different samples before and after GC correction. This figure demonstrates that GC correction significantly (p<2.2e-16; Wilcoxon signed rank test with continuity correction using the absolute values of the Pearson’s correlation value) reduced the Pearson’s correlation between copy ratios of different samples. (**C**) The theoretical coefficient of variations (TCVs) and the actual coefficient of variations (ACVs) of copy ratios before and after GC correction. The ACVs after GC correction are much closer to the lower bounds of TCVs than the ACVs without GC correction. ACVs after GC correction are only 0.0015(0.00028-0.0057) larger than the lower bounds of TCVs, implying that our GC correction approximately removed all the systematic errors in the copy ratios, including that caused by GC-biases.

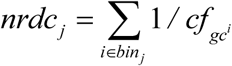

in which i represents the ID of reads whose start position located in the jth bin.

### 2.3 Control RDC construction and copy ratio calculation

To reduce systematic bias caused by factors other than GC-content, we used the copy ratio of test sample NRDC to the control reference NRDC (CNRDC) for further analysis. CNRDC is calculated by the following procedures: Firstly, to eliminate the influence of variation in sequencing data volumes, all the samples including test samples and control samples should be performed with centralization of NRDC by dividing the median NRDC of all genome bins. Secondly, to reduce the random fluctuation in the CNRDC and thus reduce the fluctuation in the final copy ratio, we construct CNRDC by averaging the NRDCs at each bin across the control samples. After obtaining the CNRDC, we can get the corrected copy ratio (CCR) for the test samples by

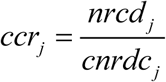

in which *ccr*_*j*_ is the corrected copy ratio of the jth bin, *nrdc*_*j*_is the NRDC of the jth bin in the test sample, and *cnrdc*_*j*_ is the CNRDC of the jth bin. Additionally, to avoid the frequent germline CNVs, outlying bins with CNRDC < *m*_*CNRDC*_ *0.7 or CNRDC > *m*_*CNRDC*_ *1.4 were removed, in which *m*_*CNRDC*_ is the median value of CNRDC of all autosomal bins.

For detecting CNVs at various level of length, we performed CNV segmentation on detection bins (defined as level-2 bins), whose length (should be an integer multiple of level-1 bin length) are assigned by users. The copy ratio of each level-2 bin was obtained by averaging CCR of all level-1 bins within it.

### 2.4 Second step of GC-correction

Even with the first step of GC correction described in 2.2, we observed that the CCRs were still generally correlated with GC-content (Supplementary Fig S2). The remaining bias, though slightly increasing the variance of CCR, may produce false positives in several GC abnormal regions in the genome, e.g. chr1p and chr19. Thus a second step of GC-correction was performed to further remove the remaining GC bias, by a generalized linear regression for CCR and GC-content of level-2 bins, which can be illustrated as:

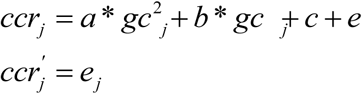

in which *a, b* and *c* are coefficients of regression, *gc* _*j*_ is GC-content of jth level-2 bin, 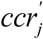 is the GC-corrected copy ratio of jth level-2 bin, and *e* _*j*_ is the residual of *ccr*_*j.*_ Then, 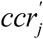 was used instead of *ccr*_*j*_ as input for CNV segmentation algorithm.

### 2.5 CNV Segmentation

To locate the breakpoints of CNVs and determine the CNV status of segments, we employed a piecewise least squares estimation model based on stepwise regression for each chromosome, which could also be described as piecewise linear fitting of ladder type and identified the breakpoints of ladders one by one. We minimized sum of squares of residuals to get the least squares estimation, as described below.

If we have obtained the sorted breakpoint set *E* ={(0,1),(*j*_2,_ *j*_2_ +1),(*j*_3,_ *j*_3_ +1),,(*j*_*k,*_ *j*_*k*_ +1),(*n,n* +1)} (1 < *j*_2_ < *j*_3_ <… < *j*_*k*_ < *n,* each breakpoint is indicated by two consecutive genomic bins and n is the total bin numbers of this chromosome) after fitting k ladders to a chromosome, we can obtain the estimation of *ccr*_*j*_ for each bin:

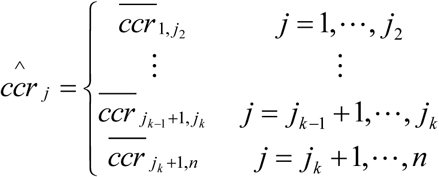

in which 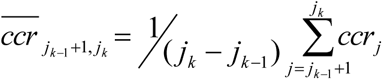, or the mean value of *ccr*_*j*_ from (*j*_*k* –1_ +1) th bin to *j*_*k*_ th bin. So the sum of squares of residuals is

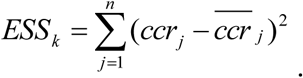

Next, we introduce the detailed steps of the piecewise linear fitting model by taking the chromosome i for example. For the jth bin of chromosome i, let 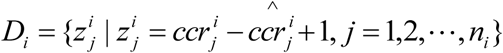 be the discriminant set for chromosome i, in which *n*_*i*_ is the total bin number.

Step 0: Get the initial value of coefficient of variation (CV). We firstly calculated CV of each 20 consecutive level-2 bins in all autosomes by

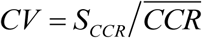

where *S*_*CCR*_ and 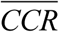 are the standard variation and the average of CCRs, respectively.. Then the initial CV (*cv*_0_) could be estimated by averaging the smallest 30% CVs of all 20 consecutive level-2 bins in the whole genome, and subsequently be used in the loop termination conditions for the piecewise linear fitting. When k=1, the breakpoint set is 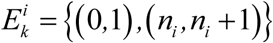, which has only two breakpoints and one CNV segment. The estimation of 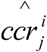 could be calculated by the mean of all 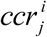 in chromosome i.

Step 1: Judge whether to terminate the loop of piecewise linear fitting for chromosome i. For discriminant set *D*_*i,*_ we can calculate the CV value of ladder k (*cv*_*k*_) by

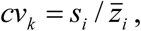

in which 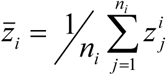 is the mean value of *D*_*i*_ set, 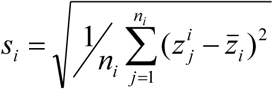 is the standard variation of *D*_*i*_ set. When k=1, if *cv*_*k*_ – *γ* \**cv*_0_ ≤ 0 (*γ* is a coefficient assigned by users, and *γ* = 1 by default) we terminate the loop for piecewise linear fitting. And when k>1, if

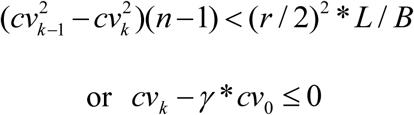

we terminate the loop, where r is a coefficient assigned by users corresponding to the lowest chimeric fraction for CNV detecting (3% by default), B is the length of level-2 bin (1M bp by default) and L is the smallest size for CNV detecting (3×b, by default). Otherwise, k=k+1, and then go to step 2.

Step 2: Add one optimal point to breakpoint set 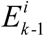 by traverse the potential new breakpoint set 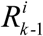. After adding one more breakpoint 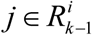 to 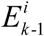, we can get the new breakpoint set 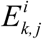 and the new sum of squares of residuals 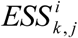. To get the optimal breakpoints for fitting the chromosome, we minimize the 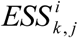 over all 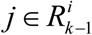:

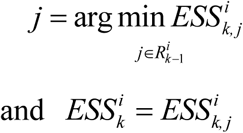

Step 3: Test whether the new breakpoint set is significant for fitting chromosome **i**. According to the stepwise regression model, if **k**>2, we should test whether including each one of the previous breakpoints in 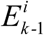 is more significant for fitting chromosome i than including the new breakpoint in step 2. If including a previous breakpoint is not more significant than including the new breakpoint, this previous breakpoint should be removed from the breakpoint set while k=k-1, until all the remained breakpoints are more significant than the new breakpoint. Then, go to step 1. The significance of including a breakpoint is assessed by calculating the decrease in ESS.

### 2.6 Significance test

For chromosome i of the test sample, we assume that 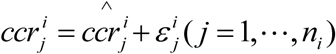 and 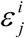 follow independent identical normal distribution 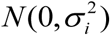. The unbiased estimation of 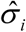 can be calculated by

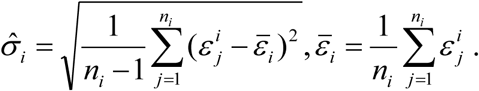

So the CCR in a normal bin should follow distribution of 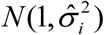. Therefore, we can inferred that average CCR of the kth segment 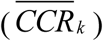 follows distribution of 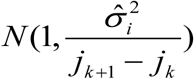 under the H0 hypothesis of 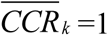, and the significance of the kth segment to reject H0 could be tested.

Because the additional deviation in the copy ratios induced by real CNVs could not be determined before going through the pipeline, the first run of step 2.5 and 2.6 was only used to define normal regions in the whole genome, and thus used to set parameters *cv*_0_ for the second run of these two steps to obtain the final segmentation results and the significance of each CNV. Before the second run of step 2.5 and 2.6, the copy ratios across the genome should be centralized again using the average copy ratio of normal bins defined by the first run. The initial coefficient of variation *cv*_0_ used in the step 0 of 2.5 in the second run is also reset by *cv* of normal bins at the first run.

## 3. Results

### 3.1 Theoretical limitation of CNV detectability in cf-DNA

The sequenced DNA fragments only took a very small part of the cf-DNA in the circulation, so the process of blood drawing, cf-DNA extracting and library construction could be regarded as random sampling of cf-DNA from an infinite population. Thus the read count examined in a certain genome bin should follow Poisson distribution. To calculate theoretical limitation of CNV detectability in cf-DNA, we firstly assessed the theoretical random fluctuation of copy ratios. The RDC in the ith bin could approximate to a random variable following Poisson distribution *P*(*λ*_*i*_) with a mathematical expectation of *λ*_*i.*_ However, *λ*_*i*_ may be different for different i because some inherent characteristics in different genomic regions could affect the read count, such as the mappability (Ha, *et al*., 2012). We have:

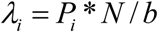

in which N is the total effective reads of this sample, b is the total number of level-2 bins in the whole genome, and *P*_*i*_ is a position specific coefficient to adjust the reads count in i th bin.

Considering X and Y are independent, the random fluctuation of copy ratios could be evaluated by

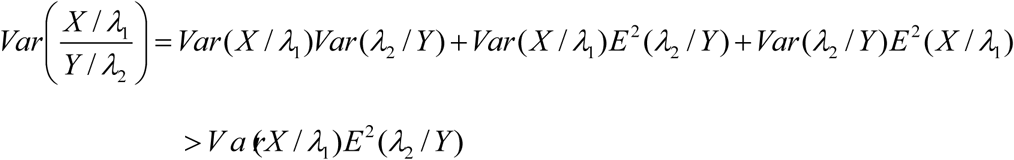

in which *λ*_1_ is mathematical expectation for test sample X and *λ*_2_ is mathematical expectation for the control reference Y.

It is easy to deduce a lower bound of CV

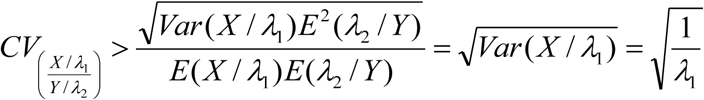

For brevity, we assume that *P*_*i*_ = 1 for all i in each sample, and use the variance of all bins in a sample as the variance of each bin. Then we can get:

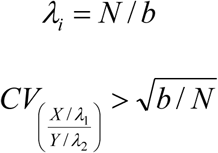

In order to evaluate the effectiveness of our GC-correction method, we applied this method to 67 normal maternal plasma samples, and compared the actual coefficient of variation (ACV) before and after GC-correction with theoretical lower limit of coefficient of variation (TCV) for the copy ratios (CR). Before GC-correction, we identified various degrees of correlation between CR and GC-content in the 67 samples, implying that different samples were affected by different levels of GC bias, even with identical number of PCR cycles in the library process. We also identified a broad range of correlation between CRs of different samples (Fig 1B), suggesting the non-independence of CR in different samples. After GC-correction, ACVs were greatly reduced, as well as the linear correlation between GC-content and CRs, and the correlation of CRs between different samples. Moreover, we approximately achieved the theoretical lower limit of TCV after GC-correction (Fig 1C). These results implied that systematic bias in CRs are mostly contributed by GC bias, which could be almost completely removed using our GC-correction method. After GC-correction, CR of different samples obeyed the assumption of independence in step 2.6. However, factors other than GC-bias that introduce systematic bias in the RDC should also affect RDC of the control samples, and thus should be normalized in the CR.

We next assessed theoretical limit of detecting power given a confidence coefficient (*p*-value). A significant CNV segmentation could be modeled as several continuous bins with deviant CRs. Approximately, we assumed that CR of single bin followed an i.i.d normal distribution, thus the detecting power could be assessed by

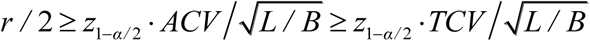

where *r* / 2 is the chimeric fraction of a heterozygous CNV, *α* is the confidence level, *z*_1–*α* / 2_ is the 1–*α* / 2 quantile of standard normal distribution, L is the length of CNV, and B is the level-2 bin size. Supplementary Figure S3 shows the smallest chimeric fraction of detectable CNV with various lengths and amount of effective reads. Given N=25M, b=3000, B=1M, L=3M, and *p*=10e-5, we could infer that *r* /2 =2.80%.

### 3.2 Simulation Study and Comparison between segmentation methods

To evaluate the performance of SCDT, we blended cf-DNA from a healthy female with DNA from tissues of aborted fetuses whose CNVs had been determined by G-banding karyotyping, to simulate cf-DNA from maternal plasma with abnormal fetus. Using this method and tissues of 11 aborted fetuses, we obtained 108 simulated samples with various mixture ratios (MR), including 3%, 5%, 8%, 10%, 15% and 20%. All the simulated samples were sequenced with 35bp single-end reads on BGIseq 500 platform. After filtering out reads with low mapping quality (<Q30) or high mismatch numbers, the average amount of effective reads for each sample is averagely 25M (8M-69M), which is applicable in current NITP test. Considering that the CNVs of aborted fetuses are different from each other, in this experiment we used the median NRDC of all the samples as the control reference for normalization.

We set the level-1 bin size to 100kb and the level-2 bin size to 1M, and then used SCDT to analyze these samples. To reduce false positives, we required a CNV segment with *p*-value smaller than 10e-5), and with copy ratio >1.01 or <0.99. Here, we defined a predicted CNV as a true positive if it overlapped with at least 50% of a spike-in CNV. Using SCDT we detected all the spike-in CNVs with chimeric fraction ≥5% (mixture ratios ≥10%) and length ≥3M and had no false positive with length ≥3M after filtering out putative germline CNVs (Fig 2). We then compared our results with the theoretically detection limitation and found that SCDT detected most of theoretically detectable CNVs, though missed some near the line of theoretically limitation (Supplementary Table S1). However, the spike-in fractions evaluated from CRs of CNVs were lower than expected (Supplementary Fig S4), probably because of failure to remove some large fragment DNA during preparation of fetal DNA.

**Figure 2.**
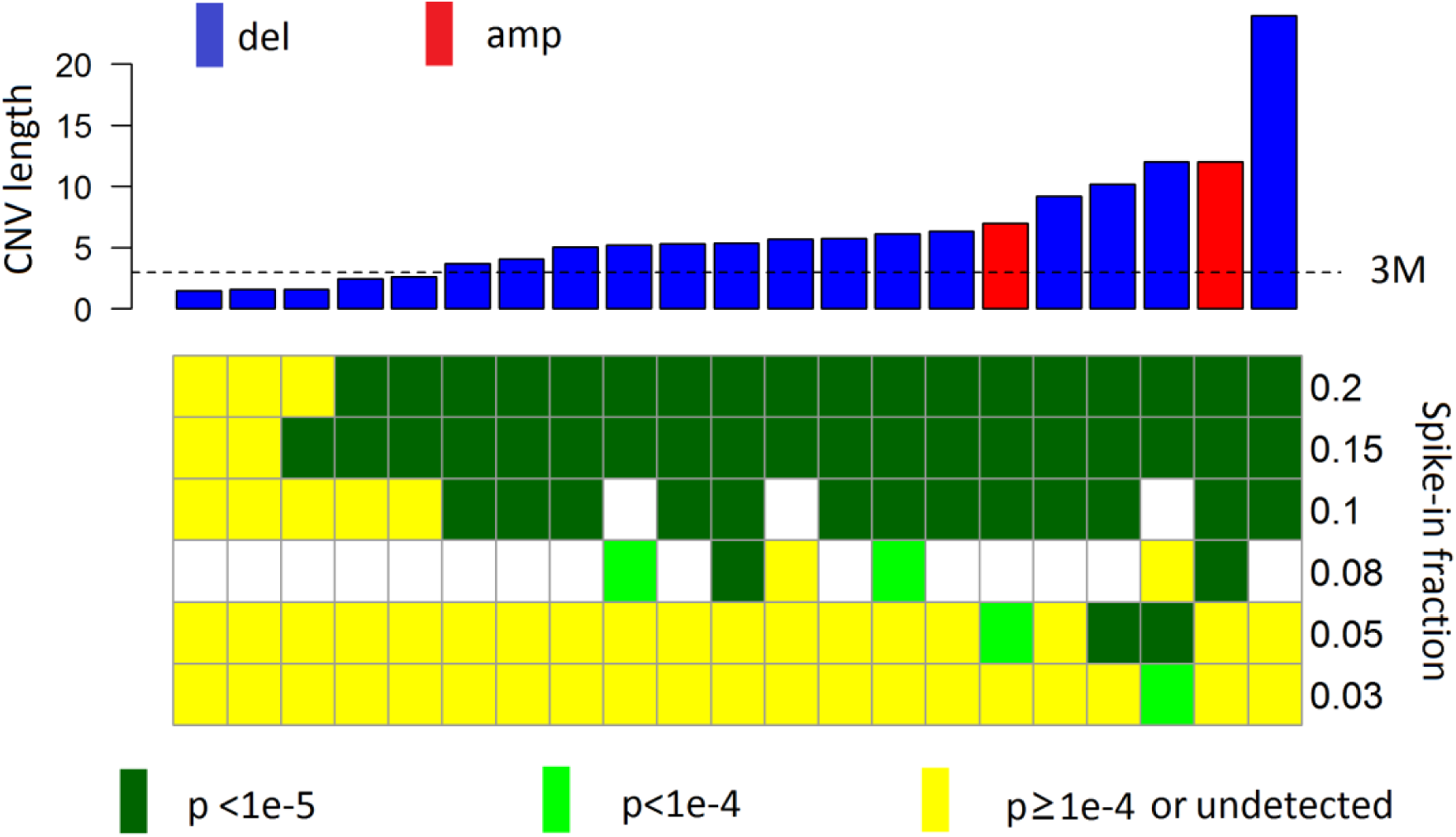
Performance of SCDS on the data of simulated maternal plasma samples. The top panel presents the lengths of spike-in amplifications (red bars) and deletions (blue bars). The lower panel presents the detecting results of spike-in CNVs, while whites blocks denote that these simulations had not been performed.

We compared the performance of SCDT on the simulated samples with the state-of-the-art CNV detection methods, including BIC-seq (Xi, *et al*., 2011), DNAcopy (Venkatraman and Olshen, 2007), Control-FREEC (Boeva, *et al*., 2012) and CNV-seq (Xie and Tammi, 2009). Considering that CBS and BIC-seq didn’t have GC correction workflow, these methods were evaluated following preprocessing by our GC-correction method. For each detector, we adjusted the parameters and cutoffs until the results achieved the fewest false positives with sensitivity of 60% (Supplementary Method). We observed that our GC-normalization step greatly improved performance of CBS and BIC-seq. For the 56 samples with theoretically detectable CNVs of size ≥ 3M, the segmentation method of SCDT had the highest sensitivity (89.29%), followed by CBS (82.14%) and BIC-seq (76.80%). When considering all the samples with the CNV size more than 3M, the segmentation method of SCDT also had higher sensitivity (59.77%) than CBS (52.87%) and BIC-seq (49.43%), while the false discovery rates for SCDT, CBS and BIC-seq were around 1.89%, 8% and 0%. We then compared the average sum of length of false positives in each sample at different level of sensitivity, and demonstrated that the segmentation method of SCDT outperformed the other methods (Fig 3).

**Figure 3.**
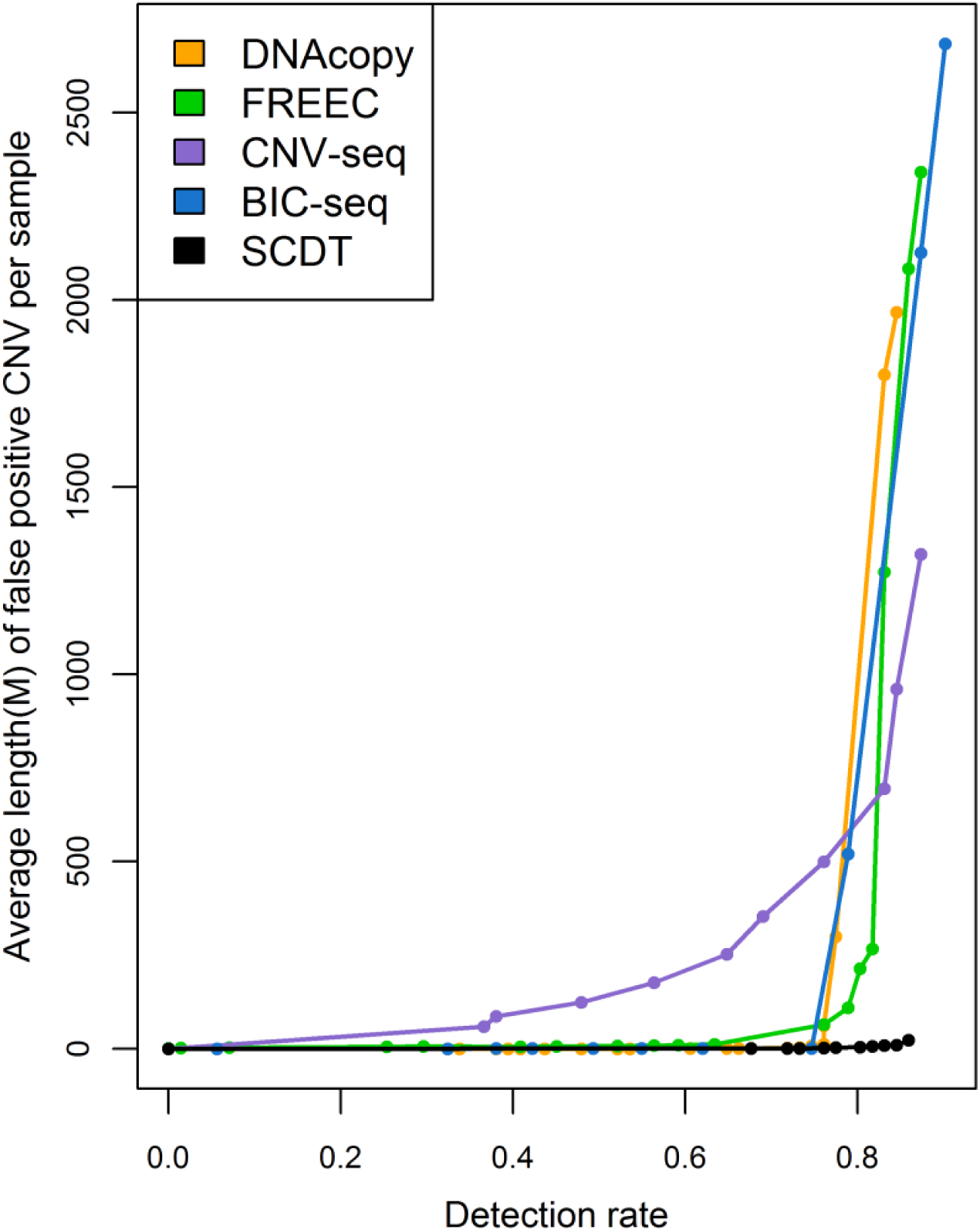
Performance of different methods on the simulated maternal plasma samples. Performances of SCDS and the state-of-the-art CNV detection methods, including BIC-seq (Xi, *et al*., 2011), DNAcopy (Venkatraman and Olshen, 2007), Control-FREEC (Boeva, *et al*., 2012) and CNV-seq (Xie and Tammi, 2009), were evaluated by the average length of all false positives in samples with certain sensitivities on the simulated maternal plasma samples (62 samples with CNV length ≥3M and spike-in fraction ≥0.05). Parameter setting of different methods was detailed in the results section.

### 3.3 Real Data Analysis3.3.1 Abnormal maternal plasma samples

To further evaluate the performance of SCDT, we applied it to cf-DNA sequencing data of real clinical samples, including maternal plasma and plasma of cancer patients. 34 maternal plasma samples carring abnormal fetal CNVs previously determined by amniotic fluid puncture and G-banding karyotyping were sequenced with average effective reads of 18M (11M - 31M). We chose another 9 normal maternal plasma samples to construct the control reference. The parameter setting was the same with the simulated data described above. We detected all confirmed CNVs in the 34 cases with only one false positive with length ≥3M, indicating high sensitivity and specificity of SCDT (Fig 4; Supplementary Table S2). We applied SCDT on another 7 cases of maternal plasma reported with abnormal genotyping by BGI NIPT workflow, but reported negative by amniocentesis and G-banding karyotyping. Interestingly, we observed chimeric CNVs in all these 7 cases, with great significance (Supplementary Fig S5). None of these women had been identified with a cancer, and inconsistence in these cases might result from chimeric placenta, abnormal hematologic clones of maternal, or false negative reports of amniocentesis.

**Figure 4.**
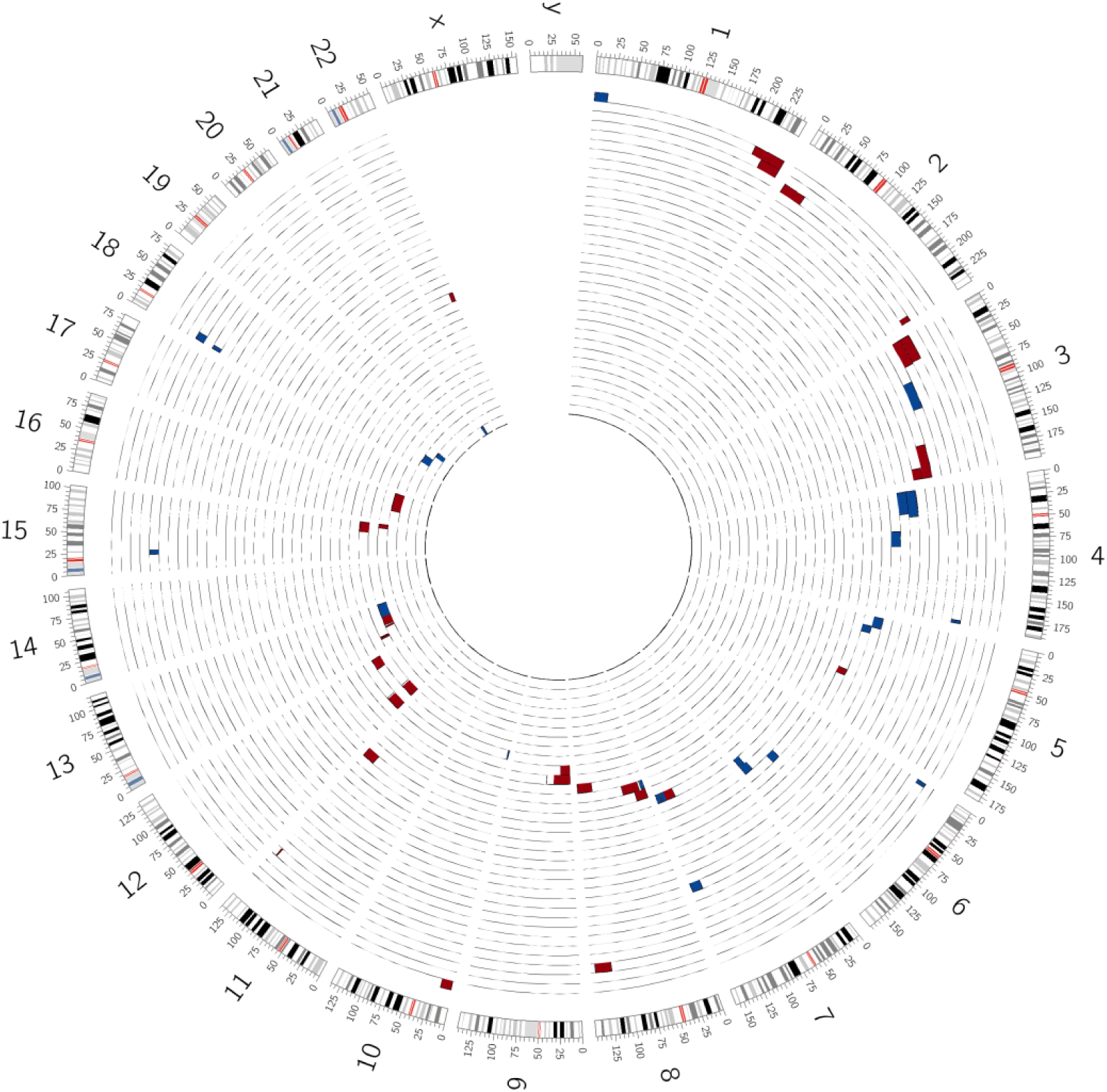
CNVs detected in the real clinical maternal plasma DNA samples carrying abnormal fetal DNA. Each sample is presented by an annular area between two adjacent circular lines. Amplifications and deletions are highlighted by red and blue, respectively.

#### 3.3.2 Target sequencing of cf-DNA for normal individuals

We then investigated CNVs in 118 cf-DNA target capture sequencing data from normal individuals. All these samples were target enriched using a panel covering 1.7 megabases before sequenced with paired-end 100bp reads on Hiseq2500 platform. To detect CNVs using off-target reads as analogue of low-depth WGS data, we filtered out reads on or near target regions to build whole genome sequencing depth for these plasma samples. The number of off-target reads is 58M averagely (27-88M). Using p<10e-5 for cut-off and filtering out a few frequent false positive regions (totally 27M) that are affected by polymorphic CNVs or harbor long centromere and telomere sequence, the false positive callings was evaluated to be 0.03% of the whole genome (Supplementary Fig S6). However, the false discovery rate should be overestimated, for some of false positives were induced by germline events.

#### 3.3.3 Target sequencing of cf-DNA for cancer patients

Finally, the same method for analyzing target-off reads was applied on additional 240 cf-DNA samples of patients with various types of cancers (Supplementary Table S3), which were captured by the same target panel used for normal individuals, with off-target reads of 45M averagely (16-170M). 10 female samples in the normal dataset were used to build the control reference. Blood cell DNA of these patients was also target sequenced as control to determine somatic point mutations. The most frequent copy number changes in 240 samples included gains of chr1q, 3q, 8q and loss of chr1p, 4, 8p, 17p, consisting with the CNV profiles in cancers (Fig 5, Supplementary Table S4). We evaluated the concentration of ct-DNA in cf-DNA using both point mutations and CNVs, and identified a high correlation between them (r=0.72, Supplementary Method). Samples detected with CNVs have significantly more point mutations and higher mutational variant allele frequency (VAF, p<1e-10) (Fig 5). Of the 107 samples that have >10% of the genome detected with CNVs, 97 (90.7%) were with average mutational VAF >2%, while in other 86 cancer samples that have ≤1% of the genome detected with CNVs,, only 6 (7.0%) were with average mutational VAF >2% and another 54 (62.8%) sample were absence of point mutations. In addition, we used VAF of point mutations to assess the copy number of CNVs, and detected several targetable agents including high amplification (CN≥7) of EGFR (n=9), MET (n=3), ERBB2 (n=5), KRAS (n=5) FGFR1 (n=1), CCND1 (n=7), CDK4 (N=4) et al (Fig 6; Supplementary Fig S7). However, high amplification (CN≥7) of some other genes known to have a role in drug resistance could also be detected in several samples, such as MYC (n=8) and MCL1(n=4), etc.

**Figure 5.**
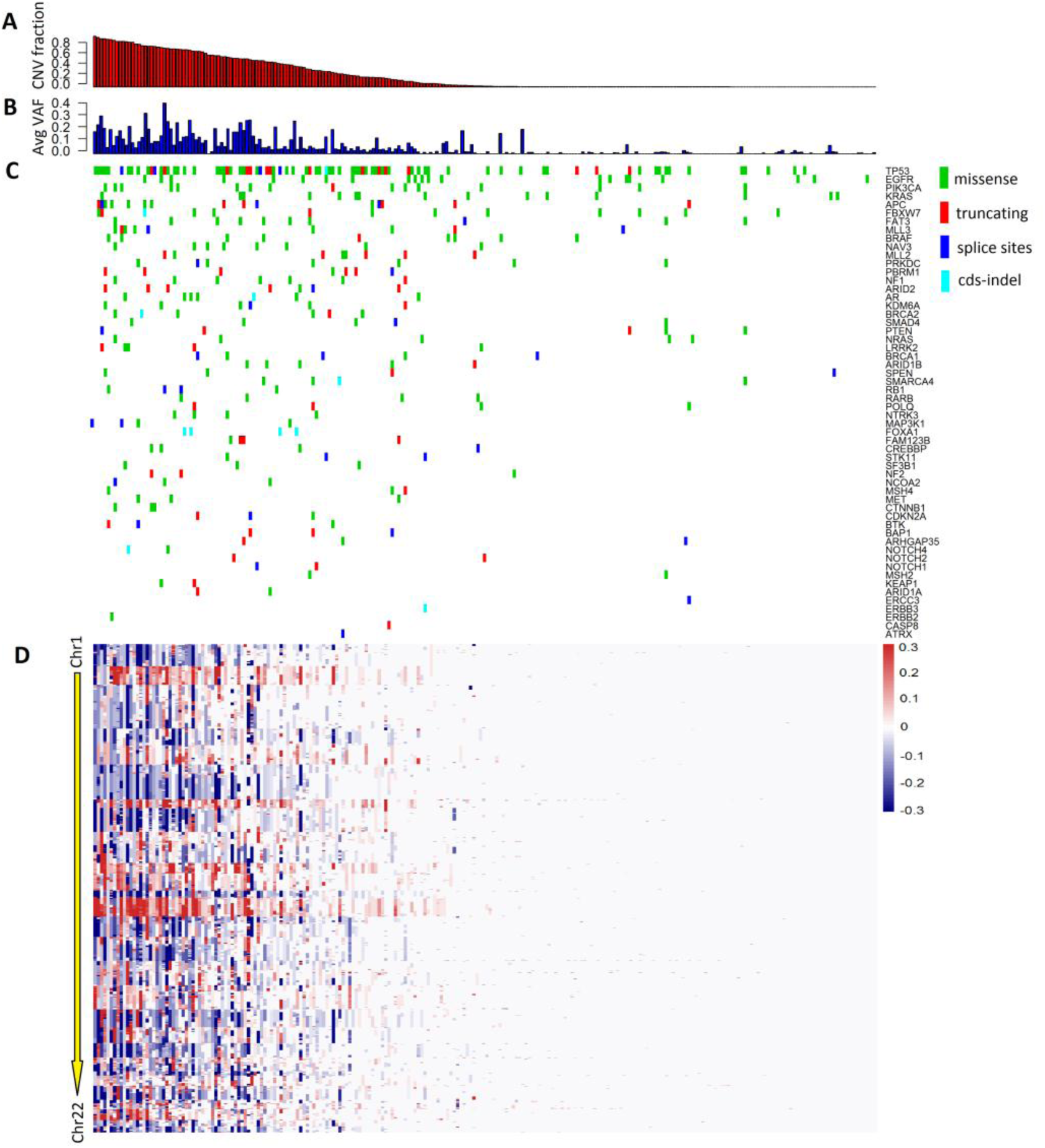
Mutations and CNVs detected in the cf-DNA of 240 cancer patients. (A) Fraction of CNV regions in the whole genome of each sample are showed by red bars. (B) Average variant allele frequencies (VAFs) of detected mutations in each sample are showed by blue bars. (C) Mutation spectrum of 240 cf-DNA samples. (D) CNV spectrum of 240 cf-DNA samples, ordered by genome positions from chromosome1 (top) to chromosome22 (bottom). Gains and losses were detected with the cutoff of p<10e-5, and highlighted by red and blue, respectively.

**Figure 6.**
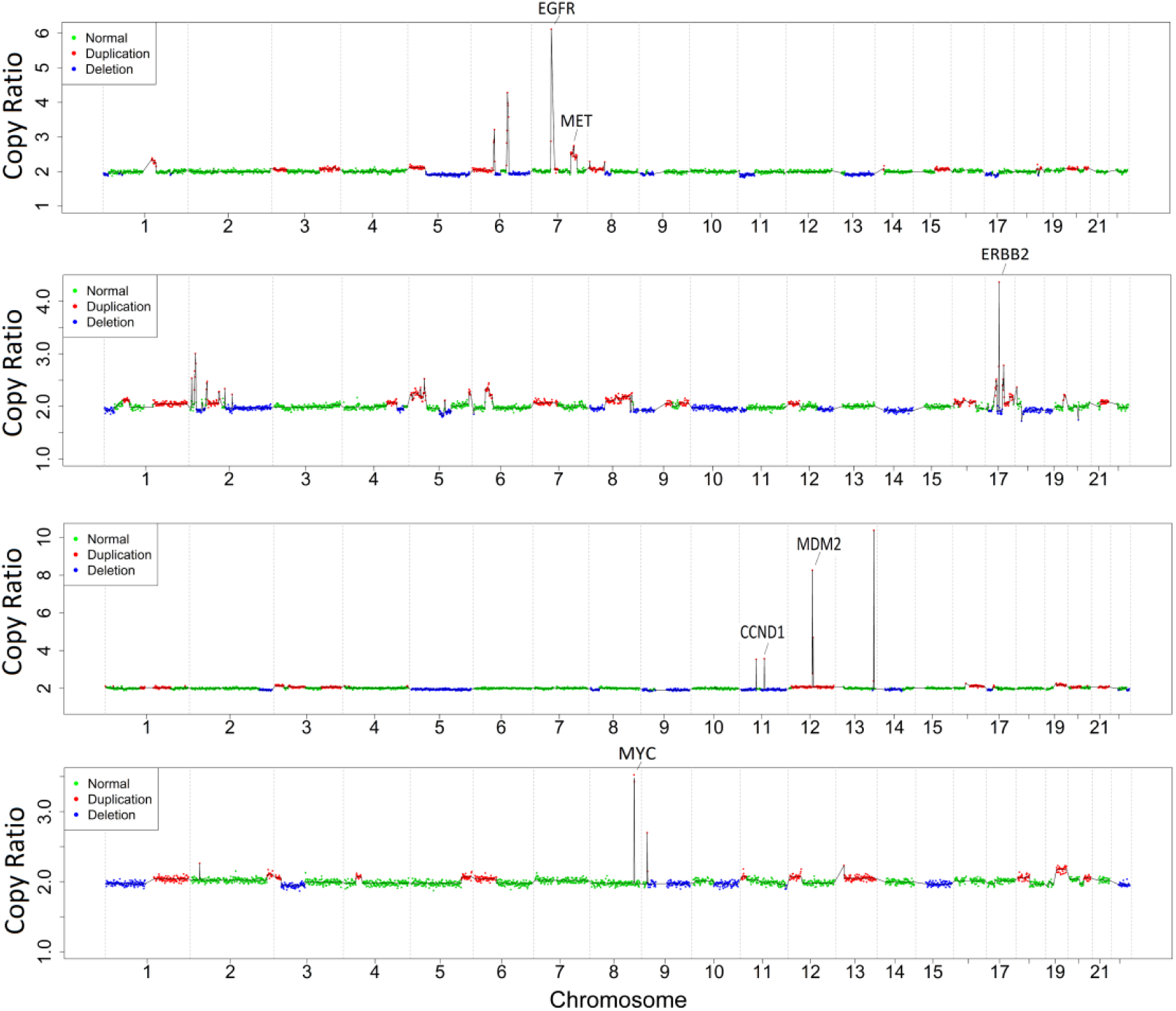
Examples of clinical relevant CNVs in cf-DNA of cancer patients. Gains and losses are highlighted by red and blue, respectively. The average VAF of mutations in four samples (from top to bottom) were 3.40%(n=11), 3.78%(n=24), 3.45%(n=5), 4.05%(n=4), respectively.

## Discussion

Deviation in the copy ratio of each genome bin is composed of at least two components, one is natural fluctuation caused by the random sampling process in blood drawing, DNA extracting, library constructing and sequencing, and the other is systematic fluctuation caused by factors such as GC non-uniform distribution or different success rate of sequencing or reads mapping in certain genome regions. According to the theoretical statistic rules, the natural fluctuation would result in poisson distribution of the final read count per genomic region, and thus can be exactly evaluated. We showed that the limit detection ability for a CNV with certain chimeric fraction and length only depends on the read count per bin, according to the theoretical statistic rules. In this study we introduced a GC correction approach to remove almost all the deviation in copy ratios contributed by non-random factors in cf-DNA sequencing data. However, as FP rate particularly concern clinical application in large scale population screening, and even a low FP rate could produce considerable FP cases, avoiding FPs of biological sources, including germline CNVs, placental mosaicism and maternal abnormality, are noteworthy in addition to reducing FPs of technical source. Several studies have identified CNVs of hemopoietic origin in blood cell-DNA in about 1-3% of normal individuals (Jacobs, *et al*., 2012; Laurie, *et al*., 2012), while they are reasonable to also present in cf-DNA, which is mainly derived from blood cell DNA (Sun, *et al*., 2015).

Cf-DNA sequencing brought immense opportunities for molecular diagnosis in clinical settings, especially in the field of cancer. However, while point mutations and methylation of cf-DNA have been widely used as biomarkers for cancer screening, early detection, treatment guiding, and disease monitoring, the cf-DNA copy-number signatures, another important kind of genomic aberrations and targets of a handful of drugs, is still seldom mentioned and evaluated in studies, partially because of the low ct-DNA fraction and technique difficulty in overcoming the low signal to noise ratio for CNV detection. Another concern is that quantifying low chimerical CNVs using WGS data requires big data size and increases the cost. However, we demonstrated that using target-off reads, the by-product of target sequencing, was a feasible way to profile somatic CNVs in cf-DNA. In addition to providing information in guiding therapeutic decisions, interrogating cf-DNA CNVs may also help to evaluate the therapy efficiency and monitor tumor recurrence. However, according to our analysis, the detection limit of chimeric ratio of CNV is inversely proportional to the square of read count, which indicates that 10 folds improvement of sensitivity requires 100 folds of sequencing data, and thus hampers the interrogating of ultra-low chimeric CNVs. Therefore point mutations may be better biomarkers for samples with low ct-DNA fraction. Moreover, as ct-DNA fraction in early stage cancers is extremely low (0.1% or less), CNVs in ct-DNA are hard to be profiled and thus not economically applicable to early cancer detection. Overall, integrating CNV analysis into liquid biopsy offers a global view of genomic aberrations, and provides more opportunity for clinical management of cancer.

To accurately find more common chromosomal abnormalities with smaller size at earlier gestational stage and cancer stage is an important goal of non-invasive clinical testing. As continuous reduction in sequencing cost, larger data size will become available soon with affordable cost. This allows for more precise diagnostics as our method is expected to perform better with increased sequencing depth. A higher coverage will allow for more stable calls and using smaller bin sizes while keeping the read depth per bin high enough to detect changes confidently.

## Acknowledgements and Funding

The research reported in this work was supported by BGI Academy of Life Sciences and BGI Institute of big data without any official funding.

## Supplementary Methods

### Parameters for BIC-seq, DNAcopy, Control-FREEC and CNV-seq

For BIC-seq, the initial bin size was set to1M and the penalty parameter was chosen as 1.5. We chose the candidate CNVs as regions with P-values less than 0.01 and logR >0.05 or <-0.05. Default parameters were used for DNAcopy. For Control-FREEC, the parameter setting include window = 1000000, degree =4, breakPointThreshold=2.0 and forceGCcontentNormalization = 1. For CNV-seq, 0.75M was chosen as the windows size and the parameter of global-normalization was used. The other parameters include minimum-windows-required = 6 and log2-threshold changing from 0.01 to 0.1 for obtaining different levels of sensitivity.

### Estimation of ct-DNA fraction in cf-DNA

Considering that homozygous deletion of long DNA fragment (>10M) was not likely to occurred in cancer genomes, so we estimate the ct-DNA fraction using the lowest copy ratio of long DNA fragment deletions (>10M) with *p*<10e-5. Assuming that µ is the lowest copy ratio of long DNA fragment deletions, we estimate the ct-DNA fraction to be (1-µ) × 2. If no long DNA fragment deletions is detected with p<10e-5, the ct-DNA fraction was estimated to be 0.

### Test dataset

The sequence data (fastq) have been deposited in the NCBI SRA database with the accession number of SRA525461.

## Supplementary Figures

**Figure S1.**
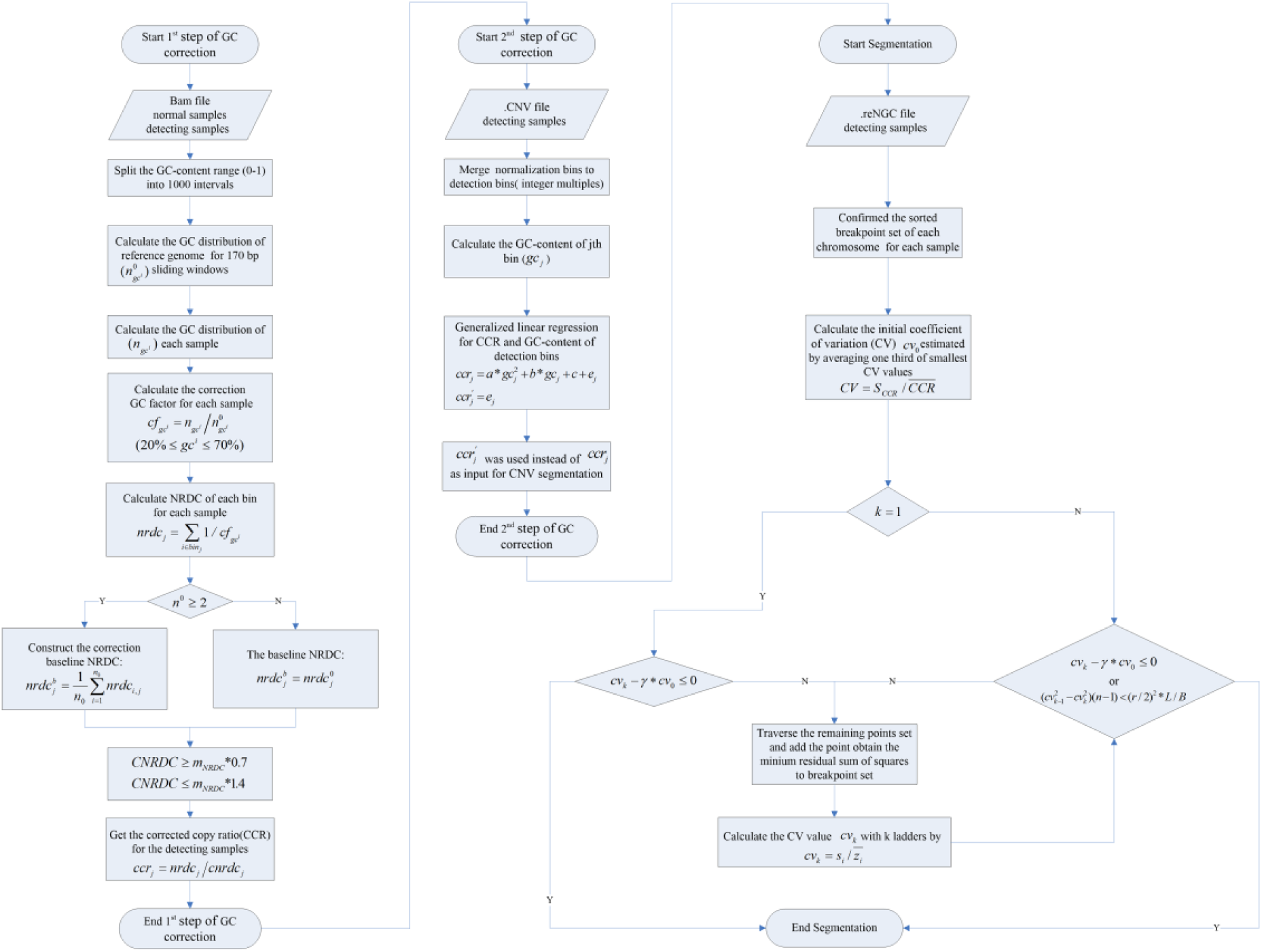
Workflow of SCDT.

**Figure S2.**
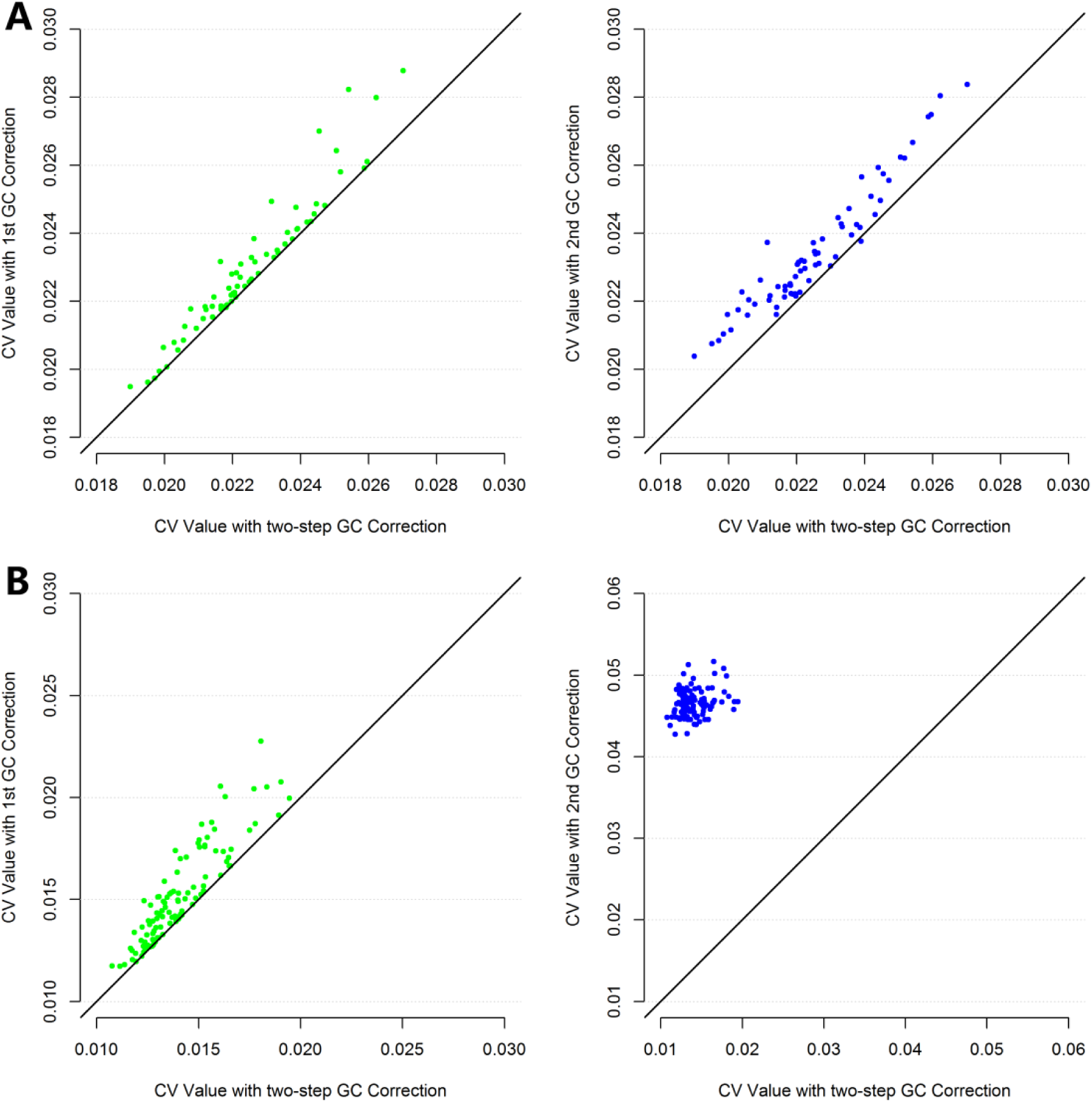
Comparison of CVs of copy ratios between two-step GC correction and one-step GC correction. CV of copy ratios after two-step GC correction (horizontal-axis) and only the first step GC correction (vertical-axis in the left panels) or only the second step GC correction (vertical-axis in the right panels) on the low-depth whole genome sequencing (WGS) data of 67 normal maternal plasma samples(A) or on the 138 target sequencing of normal samples(B).

**Figure S3.**
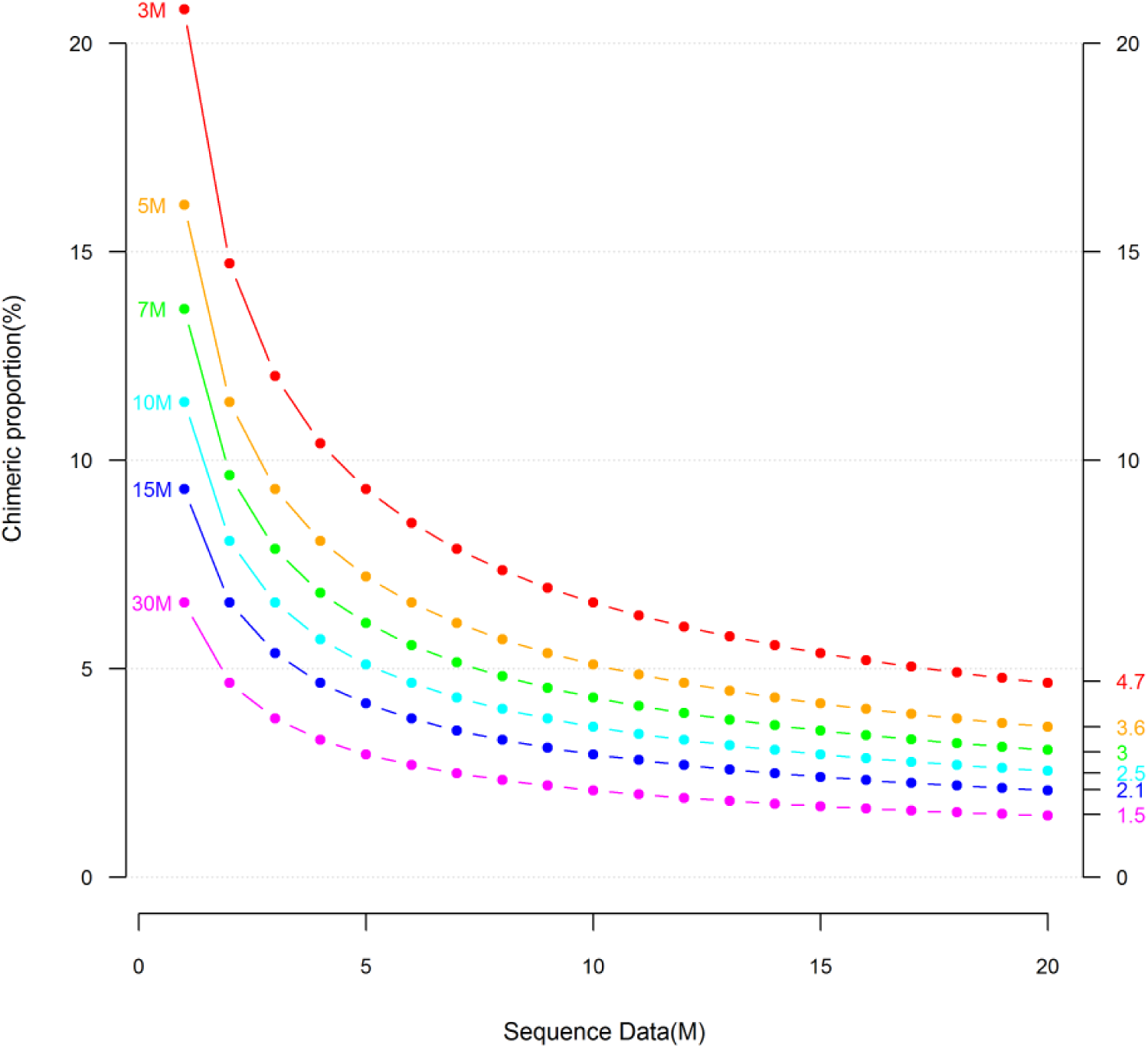
The theoretical CNV detection power determined by reads numbers, CNV length and the chimeric fraction of CNVs. The default confidence level is 0.001 and the size of test bin was set to 1M. Lines of different colors indicate the minimal detectable chimeric proportion with certain reads number and CNV length. The right vertical axis indicates the minimal detectable chimeric proportion of target CNV with 20 million reads.

**Figure S4.**
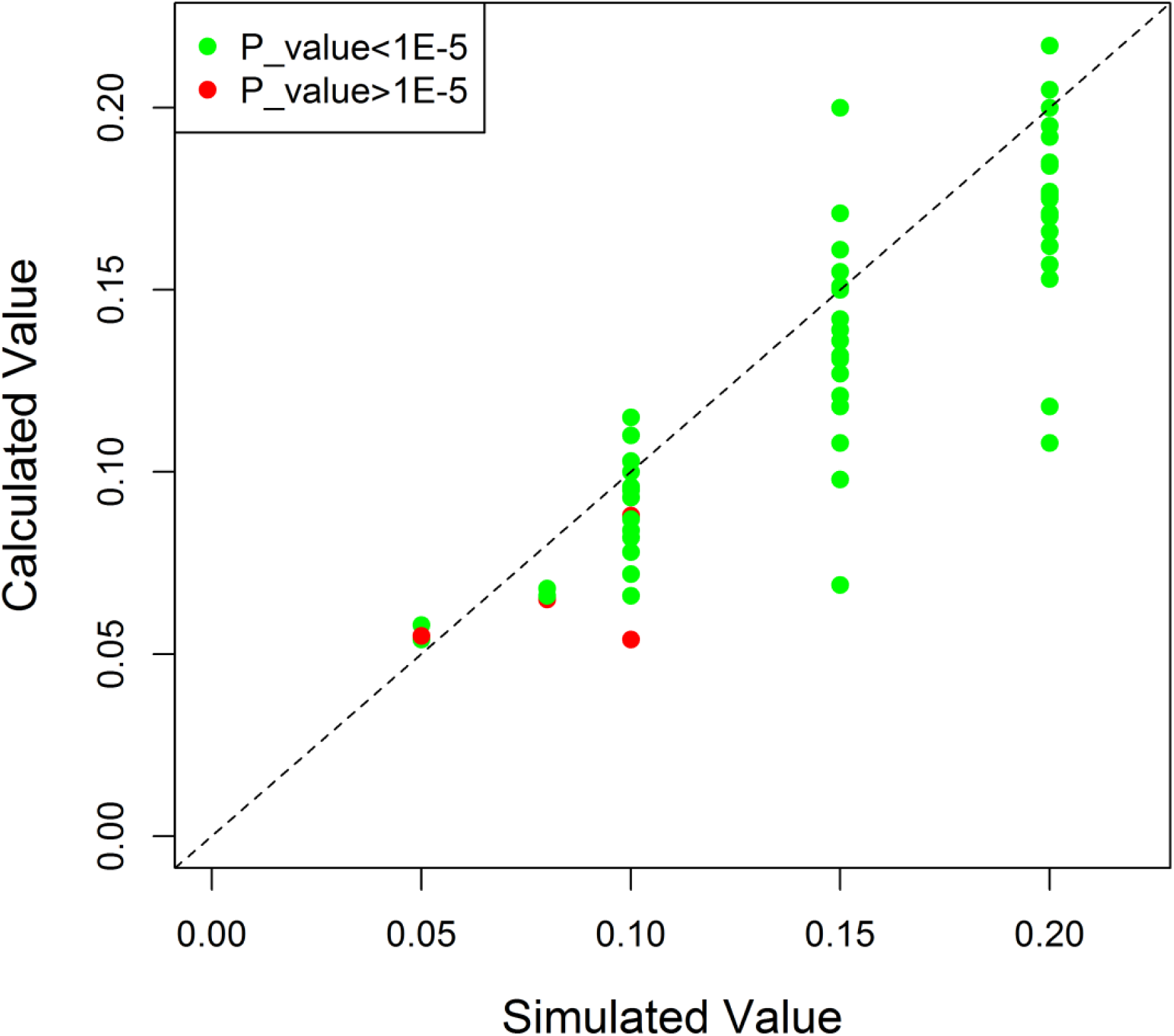
Evaluated chimeric fractions of the simulated spike-in CNVs.

**Figure S5.**
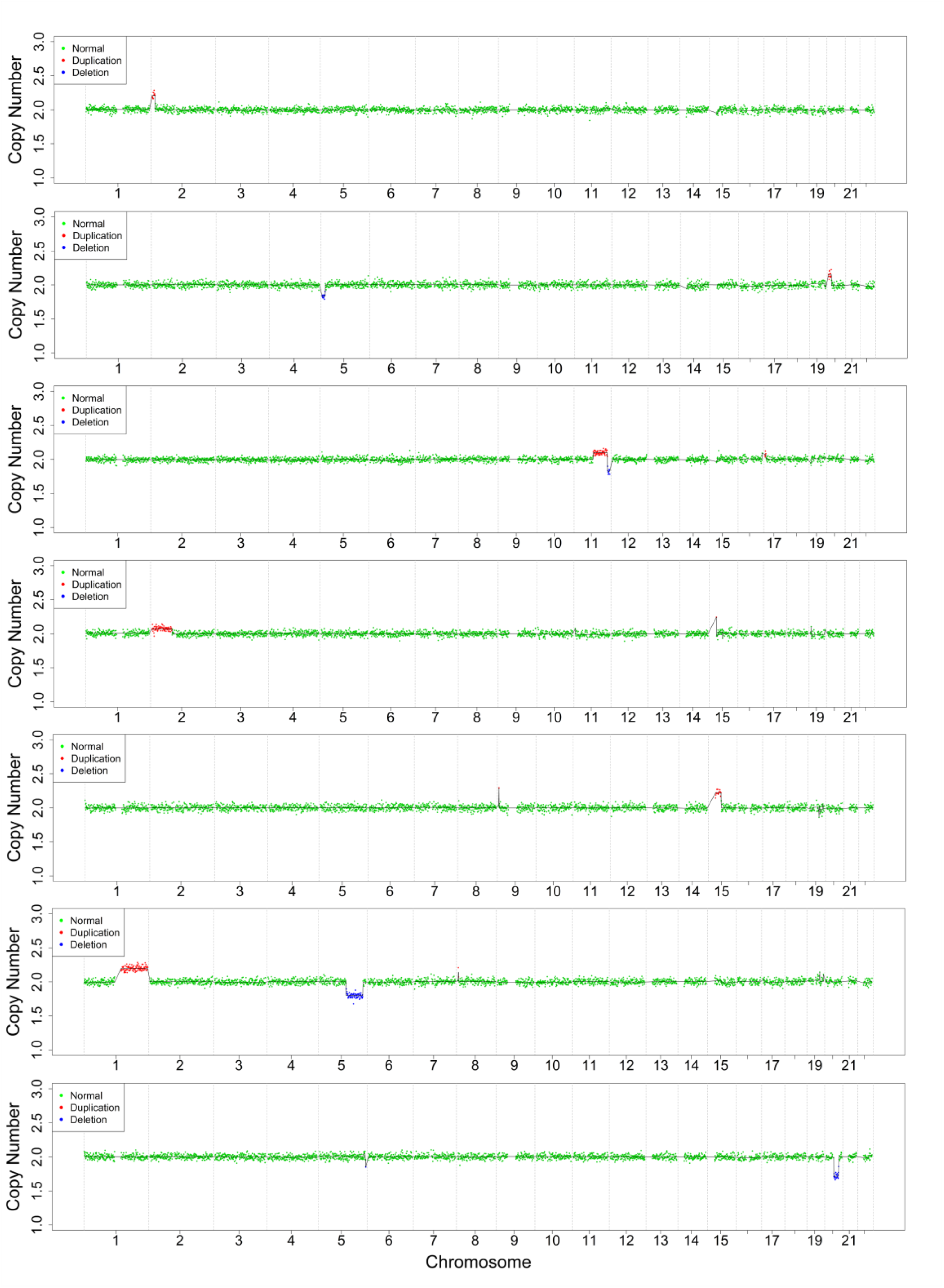
CNV profiles of 7 real maternal plasma samples which were identified to be normal by amniocentesis. Copy number gains and losses were highlighted by red and blue, respectively.

**Figure S6.**
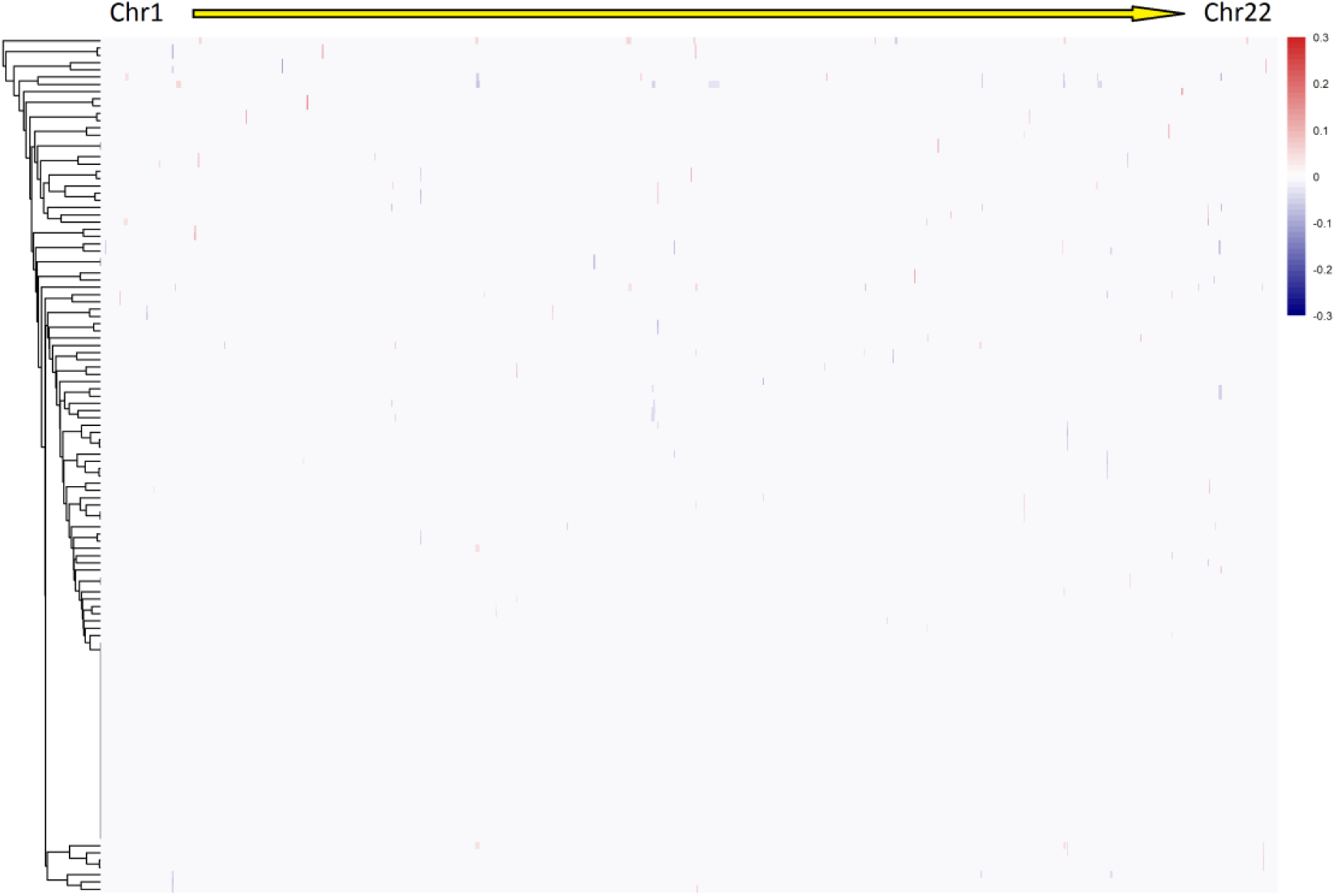
CNV spectrum in 138 target sequencing of normal cf-DNA samples, ordered by genome positions from chromosome1 (left) to chromosome22 (right). Gains and losses were detected with the cutoff of p<10e-5, and highlighted by red and blue, respectively.

**Figure S7.**
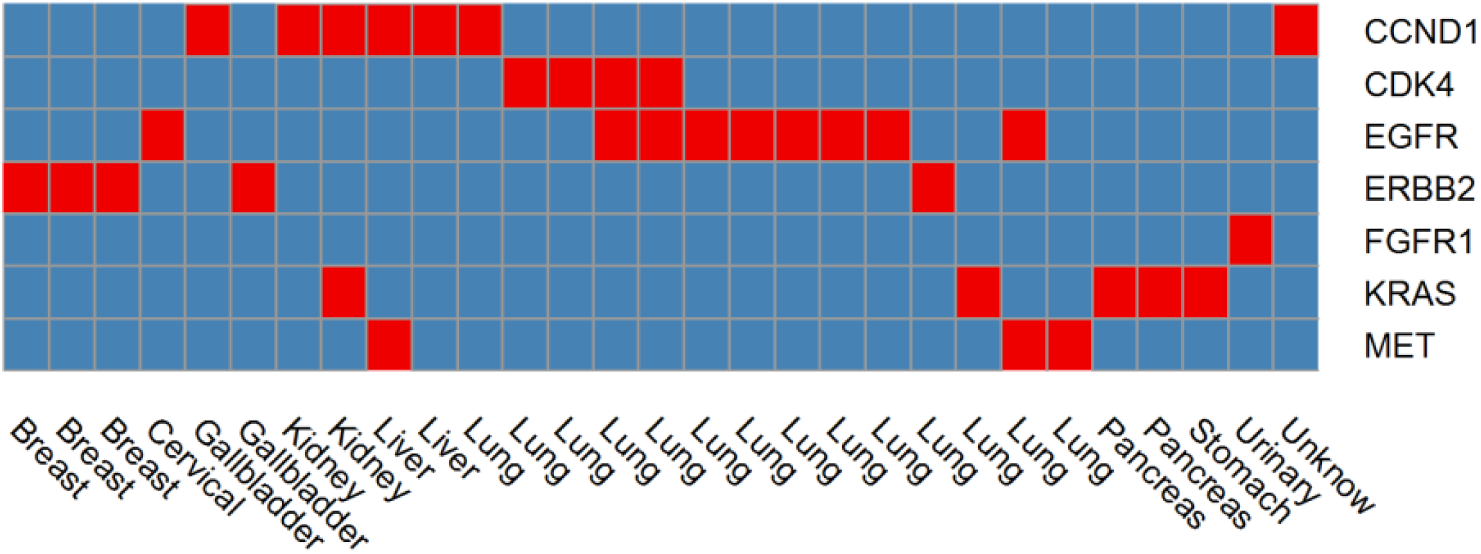
Targetable high level amplifications (CN>7) in cf-DNA of cancer patients.

## Supplementary Tables

**Supplementary table 1.**
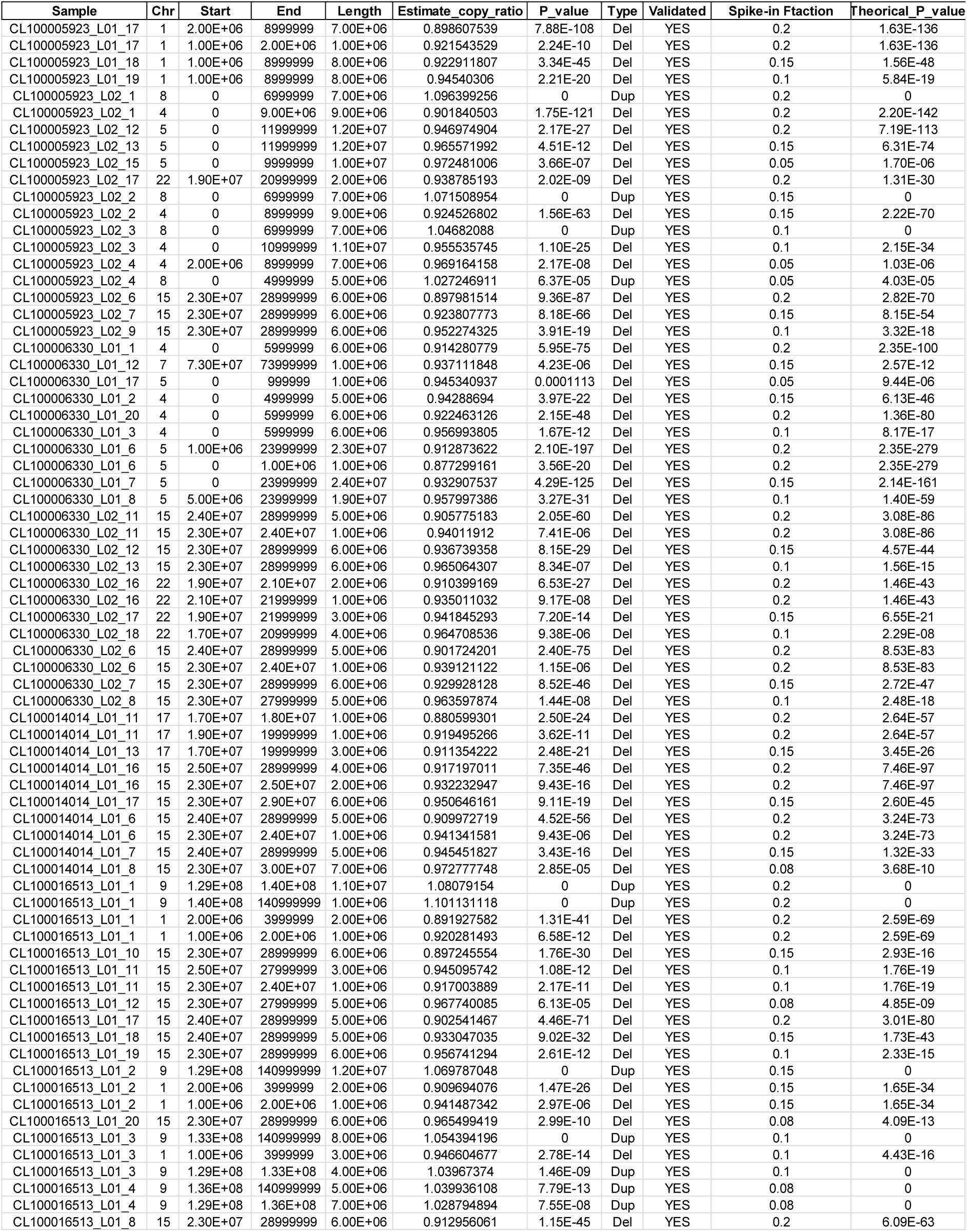

**Supplementary table 2.**

| Sample | Chr | Start | End | Length | Estimate_copy_ratio | P_value | Type | Validated |
| --- | --- | --- | --- | --- | --- | --- | --- | --- |
| CL100016513_L02_31 | 5 | 0 | 17999999 | 1.80E+07 | 0.887501211 | 1.39E-137 | Del | Yes |
| CL100016513_L02_33 | 9 | 0 | 38999999 | 3.90E+07 | 1.104866778 | 0 | Dup | Yes |
| CL100016513_L02_37 | 18 | 6.10E+07 | 77999999 | 1.70E+07 | 0.900977684 | 3.62E-106 | Del | Yes |
| CL100016513_L02_38 | 3 | 0 | 34999999 | 3.50E+07 | 1.05307488 | 0 | Dup | Yes |
| CL100016513_L02_42 | 6 | 1.03E+08 | 117999999 | 1.50E+07 | 0.938366647 | 2.52E-34 | Del | Yes |
| CL100017066_L01_12 | 3 | 1.82E+08 | 197999999 | 1.60E+07 | 1.134271942 | 0 | Dup | Yes |
| CL100017066_L01_12 | 7 | 1.43E+08 | 158999999 | 1.60E+07 | 0.867450059 | 6.58E-177 | Del | Yes |
| CL100017066_L01_13 | 3 | 0 | 36999999 | 3.70E+07 | 1.042350453 | 0 | Dup | Yes |
| CL100017066_L01_13 | 18 | 0 | 4999999 | 5.00E+06 | 0.947326212 | 2.38E-08 | Del | Yes |
| CL100017066_L01_14 | 11 | 1.10E+08 | 134999999 | 2.50E+07 | 1.04564861 | 0 | Dup | Yes |
| CL100017066_L01_14 | 6 | 1.59E+08 | 170999999 | 1.20E+07 | 0.964109986 | 4.04E-09 | Del | Yes |
| CL100017066_L01_15 | 6 | 1.62E+08 | 170999999 | 9.00E+06 | 0.862799059 | 2.12E-146 | Del | Yes |
| CL100017066_L01_15 | 6 | 1.54E+08 | 1.62E+08 | 8.00E+06 | 0.940716773 | 2.67E-21 | Del | Yes |
| CL100017066_L01_18 | 5 | 1.02E+08 | 112999999 | 1.10E+07 | 1.054948746 | 0 | Dup | Yes |
| CL100017066_L01_20 | 2 | 2.36E+08 | 242999999 | 7.00E+06 | 1.08772313 | 0 | Dup | Yes |
| CL100017066_L01_20 | 18 | 0 | 9999999 | 1.00E+07 | 0.905114039 | 1.76E-60 | Del | Yes |
| CL100017066_L01_3 | 12 | 1.07E+08 | 133999999 | 2.70E+07 | 1.06204632 | 0 | Dup | Yes |
| CL100017066_L01_3 | 10 | 0 | 2999999 | 3.00E+06 | 0.936940142 | 3.25E-13 | Del | Yes |
| CL100017066_L01_4 | 9 | 0 | 21999999 | 2.20E+07 | 1.042862869 | 0 | Dup | Yes |
| CL100017066_L01_4 | 12 | 0 | 33999999 | 3.40E+07 | 1.046307159 | 0 | Dup | Yes |
| CL100017066_L01_44 | 21 | 1.50E+07 | 23999999 | 9.00E+06 | 0.93609728 | 3.74E-25 | Del | Yes |
| CL100017066_L01_6 | 4 | 6.20E+07 | 85999999 | 2.40E+07 | 0.962856732 | 2.04E-17 | Del | Yes |
| CL100017066_L01_7 | 3 | 5.90E+07 | 97999999 | 3.90E+07 | 0.975397351 | 4.14E-08 | Del | Yes |
| CL100017066_L01_8 | 1 | 2.23E+08 | 248999999 | 2.60E+07 | 1.125025541 | 0 | Dup | Yes |
| CL100017066_L01_8 | 6 | 0 | 4999999 | 5.00E+06 | 0.883814707 | 1.66E-46 | Del | Yes |
| CL100017066_L01_9 | 3 | 1.50E+08 | 197999999 | 4.80E+07 | 1.074375309 | 0 | Dup | Yes |
| CL100017066_L02_21 | 13 | 2.00E+07 | 2.40E+07 | 4.00E+06 | 1.070766305 | 0 | Dup | Yes |
| CL100017066_L02_21 | 15 | 7.50E+07 | 102999999 | 2.80E+07 | 1.067374502 | 0 | Dup | Yes |
| CL100017066_L02_21 | 13 | 2.50E+07 | 26999999 | 2.00E+06 | 1.052683185 | 3.72E-06 | Dup | Yes |
| CL100017066_L02_22 | 15 | 9.10E+07 | 102999999 | 1.20E+07 | 1.090749056 | 0 | Dup | Yes |
| CL100017066_L02_23 | 13 | 5.30E+07 | 7.50E+07 | 2.20E+07 | 1.049337076 | 0 | Dup | Yes |
| CL100017066_L02_23 | 13 | 7.50E+07 | 7.80E+07 | 3.00E+06 | 1.101681725 | 0 | Dup | Yes |
| CL100017066_L02_23 | 13 | 7.80E+07 | 114999999 | 3.70E+07 | 0.935872676 | 2.49E-100 | Del | Yes |
| CL100017066_L02_23 | 13 | 4.50E+07 | 4.90E+07 | 4.00E+06 | 1.054928849 | 3.43E-09 | Dup | Yes |
| CL100017066_L02_24 | 5 | 1.70E+07 | 28999999 | 1.20E+07 | 0.918787306 | 2.39E-34 | Del | Yes |
| CL100017066_L02_25 | 8 | 0 | 22999999 | 2.30E+07 | 1.063801078 | 0 | Dup | Yes |
| CL100017066_L02_26 | 4 | 0 | 39999999 | 4.00E+07 | 0.954084728 | 8.67E-43 | Del | Yes |
| CL100017066_L02_27 | 8 | 1.20E+07 | 47999999 | 3.60E+07 | 1.053063259 | 0 | Dup | Yes |
| CL100017066_L02_27 | 8 | 0 | 7.00E+06 | 7.00E+06 | 0.938205434 | 4.30E-19 | Del | Yes |
| CL100017066_L02_29 | 10 | 2.00E+06 | 1.50E+07 | 1.30E+07 | 1.072197238 | 0 | Dup | Yes |
| CL100017066_L02_29 | 1 | 1.00E+06 | 16999999 | 1.60E+07 | 0.920613844 | 3.95E-81 | Del | Yes |
| CL100017066_L02_32 | 7 | 1.22E+08 | 1.42E+08 | 2.00E+07 | 1.055642174 | 0 | Dup | Yes |
| CL100017066_L02_32 | 7 | 1.42E+08 | 1.48E+08 | 6.00E+06 | 0.933183964 | 7.76E-25 | Del | Yes |
| CL100017066_L02_32 | 7 | 1.52E+08 | 158999999 | 7.00E+06 | 0.948429319 | 3.25E-16 | Del | Yes |
| CL100017066_L02_32 | 22 | 4.30E+07 | 50999999 | 8.00E+06 | 1.041045151 | 5.57E-11 | Dup | No |
| CL100017066_L02_32 | 7 | 1.50E+08 | 1.52E+08 | 2.00E+06 | 0.928259239 | 7.07E-11 | Del | Yes |
| CL100017066_L02_33 | 1 | 2.09E+08 | 248999999 | 4.00E+07 | 1.058877287 | 0 | Dup | Yes |
| CL100017066_L02_34 | 8 | 1.13E+08 | 145999999 | 3.30E+07 | 1.039174177 | 0 | Dup | Yes |
| CL100017066_L02_35 | 4 | 0 | 34999999 | 3.50E+07 | 0.946056263 | 1.77E-55 | Del | Yes |
| CL100017066_L02_36 | 18 | 2.00E+07 | 45999999 | 2.60E+07 | 0.954044289 | 1.72E-28 | Del | Yes |
| CL100017066_L02_37 | 2 | 0 | 30999999 | 3.10E+07 | 1.068014316 | 0 | Dup | Yes |
| CL100017066_L02_37 | 8 | 1.20E+08 | 1.41E+08 | 2.10E+07 | 1.050465712 | 0 | Dup | Yes |
| CL100017066_L02_38 | 16 | 3.40E+07 | 7.80E+07 | 4.40E+07 | 1.071450552 | 0 | Dup | Yes |
| CL100017066_L02_38 | 16 | 7.80E+07 | 89999999 | 1.20E+07 | 1.098995091 | 0 | Dup | Yes |
| CL100017066_L02_45 | 12 | 0 | 33999999 | 3.40E+07 | 1.054389375 | 0 | Dup | Yes |

**Supplementary table 3.**

| Sample | Cancer Type | Number of mutations | Tumor fraction estimated by SNV/indel | Tumor fraction estimated by CNV |
| --- | --- | --- | --- | --- |
| 14P1008444 | Cardia | 7 | 0.209285714 | 0.351433966 |
| 15P6653209-1 | Gallbladder | 0 | 0 | 0 |
| 15P6653177-1 | Gallbladder | 9 | 0.203888889 | 0.24462597 |
| 14P1011925-1 | Gallbladder | 17 | 0.051294118 | 0.089929658 |
| 14P1113003 | Unknow | 4 | 0.04025 | 0.038773001 |
| 15P6219946-1 | Lung | 0 | 0 | 0 |
| 15P6226246-1 | Lung | 1 | 0.012 | 0 |
| 15P6653214-1 | Lung | 0 | 0 | 0 |
| 15P6653255-1 | Lung | 1 | 0.012 | 0 |
| 15P6658474-1 | Lung | 2 | 0.0255 | 0 |
| 14P1006997 | Lung | 0 | 0 | 0 |
| 14P1114097-1 | Lung | 0 | 0 | 0 |
| 14P1114529-1 | Lung | 0 | 0 | 0 |
| 15P0201526-1 | Lung | 0 | 0 | 0 |
| 15P0201532-1 | Lung | 0 | 0 | 0 |
| 15P6219948-1 | Lung | 0 | 0 | 0 |
| 15P6226220-1 | Lung | 0 | 0 | 0 |
| 15P6651893-1 | Lung | 1 | 0.014 | 0 |
| 15P6652494-1 | Lung | 0 | 0 | 0 |
| 15P6653243-1 | Lung | 0 | 0 | 0 |
| 15P6615698-1 | Lung | 0 | 0 | 0 |
| 15P6651217-1 | Lung | 6 | 0.061666667 | 0.080435927 |
| 15P6653199-1 | Lung | 0 | 0 | 0 |
| 15P6653269-1 | Lung | 7 | 0.040285714 | 0.052360092 |
| 14P1113568 | Lung | 0 | 0 | 0.032423453 |
| 15P6658590-1 | Lung | 0 | 0 | 0.025570068 |
| 15P6658735-1 | Lung | 4 | 0.05975 | 0.071655274 |
| 15P6652529-1 | Lung | 3 | 0.035666667 | 0.065412934 |
| 14P1113521 | Lung | 0 | 0 | 0 |
| 15P6219928-1 | Lung | 13 | 0.059923077 | 0.070504555 |
| 15P6653082-1 | Lung | 5 | 0.1208 | 0.108767513 |
| 15P6652808-1 | Lung | 1 | 0.006 | 0 |
| 15P6652311-1 | Lung | 0 | 0 | 0 |
| 15P0201548-1 | Lung | 9 | 0.048666667 | 0.056340384 |
| 15P6653173-1 | Lung | 7 | 0.181 | 0.17344066 |
| 14P1112646 | Lung | 0 | 0 | 0 |
| 15P6651204-1 | Lung | 17 | 0.110705882 | 0.088645439 |
| 14P1114306-1 | Lung | 3 | 0.027333333 | 0.161816718 |
| 15P1825167-1 | Lung | 7 | 0.059714286 | 0.073946443 |
| 15P6658740-1 | Lung | 0 | 0 | 0.110454377 |
| 15P6215168-1 | Lung | 11 | 0.093636364 | 0.096164398 |
| 15P0201565-1 | Lung | 0 | 0 | 0 |
| 14P1012082 | Lung | 1 | 0.01 | 0 |
| 15P6226247-1 | Lung | 2 | 0.0175 | 0.280374985 |
| 15P6653260-1 | Lung | 5 | 0.206 | 0.303675861 |
| 15P6653227-1 | Lung | 3 | 0.068 | 0.299663748 |
| 15P6220091-1 | Lung | 11 | 0.137181818 | 0.081197413 |
| 15P0047208-1 | Lung | 6 | 0.266333333 | 0.3484067 |
| 15P6653258-1 | Lung | 3 | 0.009 | 0 |
| 15P6651928-1 | Lung | 18 | 0.226111111 | 0.266998073 |
| 15P6652785-1 | Lung | 7 | 0.086142857 | 0.107690034 |
| 15P6652362-1 | Lung | 1 | 0.143 | 0.154604273 |
| 15P6650491-1 | Lung | 13 | 0.148615385 | 0.235123776 |
| 14P1113520 | Lung | 23 | 0.169956522 | 0.123868105 |
| 15P6219987-1 | Lung | 1 | 0.01 | 0 |
| 14P1007586-1 | Lung | 0 | 0 | 0 |
| 14P1113755-1 | Lung | 0 | 0 | 0 |
| 14P1114095-1 | Lung | 0 | 0 | 0 |
| 15P6652480-1 | Lung | 0 | 0 | 0 |
| 15P6652783 | Lung | 0 | 0 | 0 |
| 15P6653171-1 | Lung | 3 | 0.007 | 0 |
| 15P6653198-1 | Lung | 0 | 0 | 0 |
| 15P6653226-1 | Lung | 0 | 0 | 0 |
| 15P6653230-1 | Lung | 2 | 0.0665 | 0.016422112 |
| 14P1012513 | Lung | 10 | 0.0702 | 0.078008495 |
| 15P6658643-1 | Lung | 2 | 0.0085 | 0 |
| 15P6658528-1 | Lung | 2 | 0.0105 | 0 |
| 15P6652977-1 | Lung | 9 | 0.071 | 0.307768202 |
| 15P6652427-1 | Lung | 2 | 0.0165 | 0 |
| 15P6653237-1 | Lung | 0 | 0 | 0 |
| 14P1114105-1 | Lung | 1 | 0.018 | 0.02133771 |
| 15P6656800-1 | Lung | 1 | 0.01 | 0 |
| 15P6658628-1 | Lung | 3 | 0.027333333 | 0.020970937 |
| 15P6651917-1 | Lung | 5 | 0.0854 | 0.127497145 |
| 14P1112647-1 | Lung | 0 | 0 | 0 |
| 14P1113491 | Lung | 1 | 0.005 | 0 |
| 14P1007923 | Lung | 0 | 0 | 0 |
| 15P6651916-1 | Lung | 0 | 0 | 0 |
| 15P6653225-1 | Lung | 3 | 0.032666667 | 0.045263244 |
| 15P6651205-1 | Lung | 1 | 0.075 | 0 |
| 14P0891664 | Lung | 6 | 0.169333333 | 0 |
| 15P6653385-1 | Lung | 1 | 0.014 | 0 |
| 15P6652980-1 | Lung | 2 | 0.058 | 0.03019133 |
| 15P6653250-1 | Lung | 7 | 0.032285714 | 0.027567631 |
| 15P6658739-1 | Lung | 1 | 0.027 | 0.027148833 |
| 15P6653239-1 | Lung | 0 | 0 | 0 |
| 14P1005328-1 | Lung | 0 | 0 | 0 |
| 15P0201529-1 | Lung | 1 | 0.005 | 0 |
| 15P0201544-1 | Lung | 0 | 0 | 0 |
| 15P6653264-1 | Lung | 2 | 0.01 | 0 |
| 14P0973622 | Lung | 2 | 0.0115 | 0 |
| 14P1011995 | Lung | 0 | 0 | 0 |
| 15P6651144-1 | Lung | 2 | 0.025 | 0.033178599 |
| 15P6653235-1 | Lung | 9 | 0.047666667 | 0.033770392 |
| 14P1114760-1 | Lung | 0 | 0 | 0 |
| 15P6653208-1 | Lung | 0 | 0 | 0 |
| 15P6651315-4 | Lung | 1 | 0.01 | 0 |
| 15P6653190-1 | Lung | 12 | 0.057416667 | 0.055379965 |
| 15P6653196-1 | Lung | 0 | 0 | 0.030212536 |
| 15P6653212-1 | Lung | 2 | 0.032 | 0.042098043 |
| 15P6658738-1 | Lung | 1 | 0.045 | 0.037710971 |
| 15P6220092-1 | Lung | 8 | 0.04775 | 0.049817947 |
| 14P1114307-1 | Lung | 0 | 0 | 0 |
| 14P1007455 | Lung | 0 | 0 | 0 |
| 15P6652773-1 | Lung | 4 | 0.0575 | 0.02997527 |
| 14P1113496 | Lung | 21 | 0.044619048 | 0.048638729 |
| 15P6653224-1 | Lung | 1 | 0.006 | 0 |
| 15P6652486-1 | Lung | 1 | 0.032 | 0.044580806 |
| 14P1113495 | Lung | 1 | 0.008 | 0 |
| 15P6219936-1 | Lung | 0 | 0 | 0 |
| 15P6654813-1 | Unknow | 1 | 0.005 | 0 |
| 15P6653144-1 | Liver | 5 | 0.0124 | 0 |
| 14P1009599 | Liver | 0 | 0 | 0 |
| 15P6653216-1 | Liver | 8 | 0.146125 | 0.196204318 |
| 15P6658494-1 | Liver | 3 | 0.088666667 | 0.092800381 |
| 14P1114322-1 | Liver | 4 | 0.23925 | 0.372007165 |
| 14P1111984 | Liver | 0 | 0 | 0.046137791 |
| 15P6653207-1 | Liver | 0 | 0 | 0 |
| 15P6653254-1 | Liver | 0 | 0 | 0 |
| 15P6653229-1 | Liver | 3 | 0.027666667 | 0.03388115 |
| 15P6657920-1 | Cervical | 1 | 0.01 | 0 |
| 15P6215252-1 | Cervical | 2 | 0.2015 | 0 |
| 15P6652285-1 | Cervical | 6 | 0.040166667 | 0.047674679 |
| 14P1114040-1 | Cervical | 0 | 0 | 0 |
| 15P6226129-1 | Cervical | 0 | 0 | 0 |
| 14P1115157-1 | Cervical | 10 | 0.0371 | 0.048833597 |
| 15P6219937-1 | Cervical | 11 | 0.034090909 | 0.061155945 |
| 15P6658736-1 | Cervical | 6 | 0.015166667 | 0.019174428 |
| 14P1012070 | Bone | 3 | 0.011333333 | 0 |
| 14P1112476 | Bone | 0 | 0 | 0 |
| 14P1012073 | Bone | 0 | 0 | 0 |
| 15P6652525-1 | Bone | 0 | 0 | 0.050822655 |
| 15P6651124-1 | Melanoma | 7 | 0.038857143 | 0.05140447 |
| 15P6653142-1 | Synovial_sarcoma | 0 | 0 | 0 |
| 15P6652777-1 | Glioblastoma | 0 | 0 | 0 |
| 14P1113000 | Thyroid | 0 | 0 | 0 |
| 14P1113002 | Colorectal | 0 | 0 | 0 |
| 14P1114106-1 | Colorectal | 0 | 0 | 0 |
| 15P6658485-1 | Colorectal | 0 | 0 | 0.269176289 |
| 15P6658716-1 | Colorectal | 11 | 0.259 | 0.174174302 |
| 15P6653204-1 | Colorectal | 5 | 0.3136 | 0.201413403 |
| 15P6515536R-1 | Colorectal | 0 | 0 | 0 |
| 14P1007519 | Colorectal | 11 | 0.210727273 | 0.211610161 |
| 15P0201540-1 | Colorectal | 3 | 0.047333333 | 0.371726724 |
| 15P6653221-1 | Colorectal | 1 | 0.006 | 0 |
| 14P1006572 | Colorectal | 3 | 0.043666667 | 0.051791102 |
| 14P0975261 | Lymphoma | 1 | 0.009 | 0 |
| 15P6653186-1 | Lymphoma | 5 | 0.1298 | 0.061800504 |
| 15P6655982-1 | Lymphoma | 36 | 0.191555556 | 0.27810369 |
| 15P6652491-1 | Ovarian | 0 | 0 | 0 |
| 15P6650887-1 | Ovarian | 1 | 0.068 | 0 |
| 15P6654500-1 | Ovarian | 0 | 0 | 0 |
| 15P6656825-1 | Ovarian | 2 | 0.02 | 0 |
| 15P6651892-1 | Ovarian | 0 | 0 | 0 |
| 15P6652359-1 | Ovarian | 4 | 0.2075 | 0.227239234 |
| 14P1004412-1 | Ovarian | 14 | 0.099357143 | 0.223643992 |
| 15P6653234-1 | Ovarian | 6 | 0.193166667 | 0.247161438 |
| 15P6653170-1 | Ovarian | 11 | 0.011818182 | 0 |
| 15P6652786-1 | Ovarian | 3 | 0.016 | 0 |
| 14P0973640 | Ovarian | 1 | 0.008 | 0 |
| 15P6656801-1 | Ovarian | 1 | 0.007 | 0 |
| 15P6653192-1 | Ovarian | 4 | 0.08775 | 0.053035986 |
| 15P6653240-1 | Ovarian | 2 | 0.005 | 0 |
| 15P6653220-1 | Ovarian | 1 | 0.005 | 0 |
| 15P6658589-1 | Urinary | 15 | 0.420133333 | 0.629339452 |
| 15P6652774-1 | Urinary | 10 | 0.0404 | 0.05919783 |
| 15P6653011-1 | Brain | 1 | 0.014 | 0 |
| 15S6651920-1 | Prostate | 6 | 0.034666667 | 0.086907028 |
| 15P6653252-1 | Prostate | 3 | 0.115666667 | 0.090258107 |
| 15P6651145-1 | Prostate | 5 | 0.223 | 0.253668689 |
| 15P6652775-1 | Prostate | 5 | 0.2786 | 0.615161866 |
| 15P6653178-1 | Prostate | 8 | 0.136625 | 0.212796787 |
| 15P6653248-1 | Prostate | 2 | 0.0065 | 0.033942454 |
| 15P6653222-1 | Prostate | 6 | 0.015666667 | 0.028462785 |
| 15P6656542-1 | Sarcoma | 2 | 0.2815 | 0.102729947 |
| 14P1113655 | Sarcoma | 0 | 0 | 0 |
| 15P6226230-1 | Breast | 0 | 0 | 0 |
| 15P6652157-1 | Breast | 5 | 0.0378 | 0.051497365 |
| 15P6651349-1 | Breast | 3 | 0.076333333 | 0.109239902 |
| 15P6653217-1 | Breast | 0 | 0 | 0 |
| 15P6653201-1 | Breast | 6 | 0.122833333 | 0.069904242 |
| 14P0973390 | Breast | 3 | 0.179666667 | 0.229327757 |
| 15P6653206-1 | Breast | 2 | 0.183 | 0.339381076 |
| 14P0951863 | Breast | 26 | 0.098884615 | 0.510480856 |
| 15P6220005-1 | Breast | 1 | 0.005 | 0 |
| 14P1005818-1 | Breast | 2 | 0.101 | 0.260254934 |
| 14P1005818-2 | Breast | 2 | 0.1915 | 0.475762298 |
| 15P6653193-1 | Breast | 6 | 0.024333333 | 0.030172081 |
| 15P6651130-1 | Breast | 0 | 0 | 0 |
| 15P0201525-1 | Breast | 0 | 0 | 0 |
| 14P1113686 | Breast | 3 | 0.013666667 | 0 |
| 15P6653218-1 | Breast | 4 | 0.2715 | 0.272666702 |
| 15P6658732-1 | Kidney | 0 | 0 | 0 |
| 15P6653189-1 | Kidney | 10 | 0.116 | 0.139971233 |
| 15P6651925-1 | Kidney | 0 | 0 | 0 |
| 15P6653185-1 | Kidney | 24 | 0.034541667 | 0.043863828 |
| 14P1007957-1 | Esophagus | 0 | 0 | 0 |
| 15P6658733-1 | Esophagus | 0 | 0 | 0 |
| 14P1010179 | Esophagus | 13 | 0.091692308 | 0.148532209 |
| 14P1112999 | Esophagus | 3 | 0.015666667 | 0.037563001 |
| 15P6652787-1 | Odiduct | 5 | 0.138 | 0.088576241 |
| 15P6653228-1 | Head&Neck | 0 | 0 | 0 |
| 15P6652981-1 | Head&Neck | 5 | 0.0334 | 0.036663166 |
| 15P6654180-1 | Head&Neck | 6 | 0.14 | 0.343960522 |
| 15P6653191-1 | Head&Neck | 3 | 0.086333333 | 0.13298039 |
| 14P1114363-1 | Unknow | 15 | 0.1008 | 0.170220157 |
| 14P1009600 | Unknow | 10 | 0.2589 | 0.221831563 |
| 14P1011058 | Unknow | 2 | 0.0095 | 0 |
| 14P1114098-1 | Unknow | 7 | 0.029142857 | 0 |
| 14P1114373-1 | Unknow | 1 | 0.017 | 0 |
| 15P6651951-1 | Stomach | 0 | 0 | 0 |
| 15P6652784-1 | Stomach | 52 | 0.144423077 | 0.294842894 |
| 15P6653261-1 | Stomach | 0 | 0 | 0 |
| 15P6653197-1 | Stomach | 0 | 0 | 0 |
| 15P6653205-1 | Stomach | 0 | 0 | 0 |
| 14P1012171 | Stomach | 0 | 0 | 0.04230244 |
| 14P1006996-1 | Stomach | 10 | 0.1993 | 0.378794082 |
| 15P6653172-1 | Stomach | 10 | 0.0269 | 0 |
| 14P1008017-1 | Stomach | 0 | 0 | 0 |
| 15P6653195-1 | Stomach | 8 | 0.0365 | 0.063398046 |
| 14P1007958-1 | Stomach | 0 | 0 | 0.042912187 |
| 15P1875721-1 | GIST | 0 | 0 | 0 |
| 15P6651691-1 | Unknow | 1 | 0.009 | 0.050555753 |
| 15P6653200-1 | Fibrosarcoma | 6 | 0.062166667 | 0.091158194 |
| 15P6653194-1 | SIST | 0 | 0 | 0 |
| 15P6653187-1 | Thymic | 6 | 0.275 | 0.26576364 |
| 14P1113517 | Pancrease | 1 | 0.006 | 0 |
| 15P6653979-1 | Pancrease | 2 | 0.1355 | 0.095874427 |
| 15P6652481-1 | Pancrease | 0 | 0 | 0 |
| 15P6658045-1 | Pancrease | 8 | 0.336625 | 0.322894514 |
| 15P6653247-1 | Pancrease | 13 | 0.011692308 | 0 |
| 14P1112533 | Pancrease | 0 | 0 | 0 |
| 14P1011866 | Pancrease | 2 | 0.0115 | 0 |
| 15P6653249-1 | Pancrease | 1 | 0.1 | 0.138650611 |
| 15P6226233-1 | Pancrease | 7 | 0.022428571 | 0.037242546 |
| 15P6653259-1 | Pancrease | 4 | 0.0405 | 0.028335921 |
| 15P6226237-1 | Pancrease | 5 | 0.1994 | 0.156404556 |
| 15P6650388-1 | Pancrease | 1 | 0.016 | 0 |
| 15P6653210-1 | Endometrium | 20 | 0.058 | 0 |
| 14P1112320 | Trophoblastic | 1 | 0.075 | 0 |

